# Temporal associations of B and T cell immunity with robust vaccine responsiveness in a 16-week interval BNT162b2 regimen

**DOI:** 10.1101/2021.12.18.473317

**Authors:** Manon Nayrac, Mathieu Dubé, Gérémy Sannier, Alexandre Nicolas, Lorie Marchitto, Olivier Tastet, Alexandra Tauzin, Nathalie Brassard, Guillaume Beaudoin-Bussières, Dani Vézina, Shang Yu Gong, Mehdi Benlarbi, Romain Gasser, Annemarie Laumaea, Catherine Bourassa, Gabrielle Gendron-Lepage, Halima Medjahed, Guillaume Goyette, Gloria-Gabrielle Ortega-Delgado, Mélanie Laporte, Julia Niessl, Laurie Gokool, Chantal Morrisseau, Pascale Arlotto, Jonathan Richard, Cécile Tremblay, Valérie Martel-Laferrière, Andrés Finzi, Daniel E. Kaufmann

## Abstract

Spacing of the BNT162b2 mRNA doses beyond 3 weeks raised concerns about vaccine efficacy. We longitudinally analyzed B cell, T cell and humoral responses to two BNT162b2 mRNA doses administered 16 weeks apart in 53 SARS-CoV-2 naïve and previously-infected donors. This regimen elicited robust RBD-specific B cell responses whose kinetics differed between cohorts, the second dose leading to increased magnitude in naïve participants only. While boosting did not increase magnitude of CD4^+^ T cell responses further compared to the first dose, unsupervised clustering analyses of single-cell features revealed phenotypic and functional shifts over time and between cohorts. Integrated analysis showed longitudinal immune component-specific associations, with early Thelper responses post-first dose correlating with B cell responses after the second dose, and memory Thelper generated between doses correlating with CD8 T cell responses after boosting. Therefore, boosting elicits a robust cellular recall response after the 16-week interval, indicating functional immune memory.

## INTRODUCTION

The coronavirus disease 19 (COVID-19) pandemic caused a race for the elaboration and deployment of prophylactic vaccines against SARS-CoV-2 (Krammer, 2020), including vaccines based on mRNA-based technologies that have shown clear efficacy (Baden et al., 2021; Dickerman et al., 2021; Skowronski and De Serres, 2021; Thomas et al., 2021). These mRNA vaccines target the trimeric Spike (S) glycoprotein that facilitates SARS-CoV-2 entry into host cells via its receptor-binding domain (RBD) (Hoffmann et al., 2020; Walls et al., 2020). Antibody responses are associated with protection for most licensed vaccines and the generation of Spike-specific antibodies, particularly of neutralizing RBD-specific antibodies, is considered critical for SARS-CoV-2 vaccine efficacy. Protective antibody responses are being identified (Earle et al., 2021; Gilbert et al., 2021) but there is a need for a better understanding of B cell memory responses in the context of different vaccine modalities. CD4^+^ T cell help is critical for development and maintenance of antibody immunity. SARS-CoV-2-specific CD4^+^ and CD8^+^ T cells may contribute to recovery from COVID-19 (Bange et al., 2021; Wurm et al., 2020). mRNA vaccines elicit CD4^+^ T cell responses (Anderson et al., 2020; Lederer et al., 2020; Painter et al., 2021; Prendecki et al., 2021; Sahin et al., 2020) that are likely important determinants of vaccine efficacy. CD4^+^ T subsets include T follicular helper (Tfh) cells that are critical for the expansion, affinity maturation and memory development of B cells (Crotty, 2019), and Th1 cells, which foster development of CD8^+^ T cell memory (Laidlaw et al., 2016). However, T cell subsets show important heterogeneity and plasticity, better fitting with spectra of phenotypes and functions than fully distinct populations (O’Shea and Paul, 2010). Unequivocal lineage characterization is therefore challenging, and unsupervised clustering analytical approaches are increasingly used to identify T cell subsets more specifically associated with immunological outcomes (Apostolidis et al., 2021; Maucourant et al., 2020).

The standard BNT162b2 immunization regimen recommends a 21-day interval between vaccine doses, and inoculation of two doses irrespective of prior SARS-CoV-2 infection status. However, the optimal interval has not been determined in controlled trials. In the context of vaccine scarcity and given the significant protection already conferred by the first dose in non-high-risk populations (Baden et al., 2021; Polack et al., 2020; Skowronski and De Serres, 2021), some public health agencies implemented schedules with longer intervals between doses to rapidly extend population coverage (Paltiel et al., 2021; Tuite et al., 2021), and recommended a single dose for previously-infected immunocompetent people. Longer delays between doses also frequently occur in real-life settings. While such strategies generated concerns given uncertain immunogenicity, a longer period of partial vulnerability to infection and a hypothetical risk of escape mutant selection, epidemiological evidence supports this approach as a valid alternative in lower-risk populations (Carazo et al., 2021; Skowronski et al., 2021) in which robust T cell and antibody responses are observed after a single dose (Tauzin et al., 2021b), and stronger and broader antibody immunity induced after the second dose (Grunau et al., 2021; Tauzin et al., 2021a; Payne et al., 2021). While significant progress has been made in the understanding of the kinetics of B and T cell responses in short interval mRNA vaccine schedules (Goel et al., 2021; Painter et al., 2021; Zollner et al., 2021), the immunological implications of widely-spaced vaccination regimens remain poorly known.

Here, we define the trajectories, differentiation state and interplay of vaccine-induced Spike-specific B cells, CD4^+^ T cells, CD8^+^ T cells and antibody responses in SARS-CoV-2 naïve or previously-infected individuals who received two mRNA vaccine doses administered 16 weeks apart, and in a third group of previously-infected individuals who received a single vaccine dose.

## RESULTS

### Study participants

We evaluated immune responses in three cohorts of health care workers (HCW) (Figure 1A): 26 SARS-CoV-2 naïve and 15 SARS-CoV-2 previously-infected (PI) donors who received a two-dose BNT162b2 regimen spaced by 16 weeks; and 12 PI who received a single dose. Blood samples were collected at 5 time points: at baseline (V0); 3 weeks after the first dose (V1); 12 weeks after the first dose (V2); 3 weeks after the second dose for participants receiving two doses (V3), or 19 weeks after the first dose for the single-dose PI participants (V3’); and 16 weeks after the second dose (V4). Clinical characteristics (Table 1) did not statistically differ between cohorts, except for the numbers of days between V0 and the first dose and for time between the first dose and V2.

**Figure 1.**
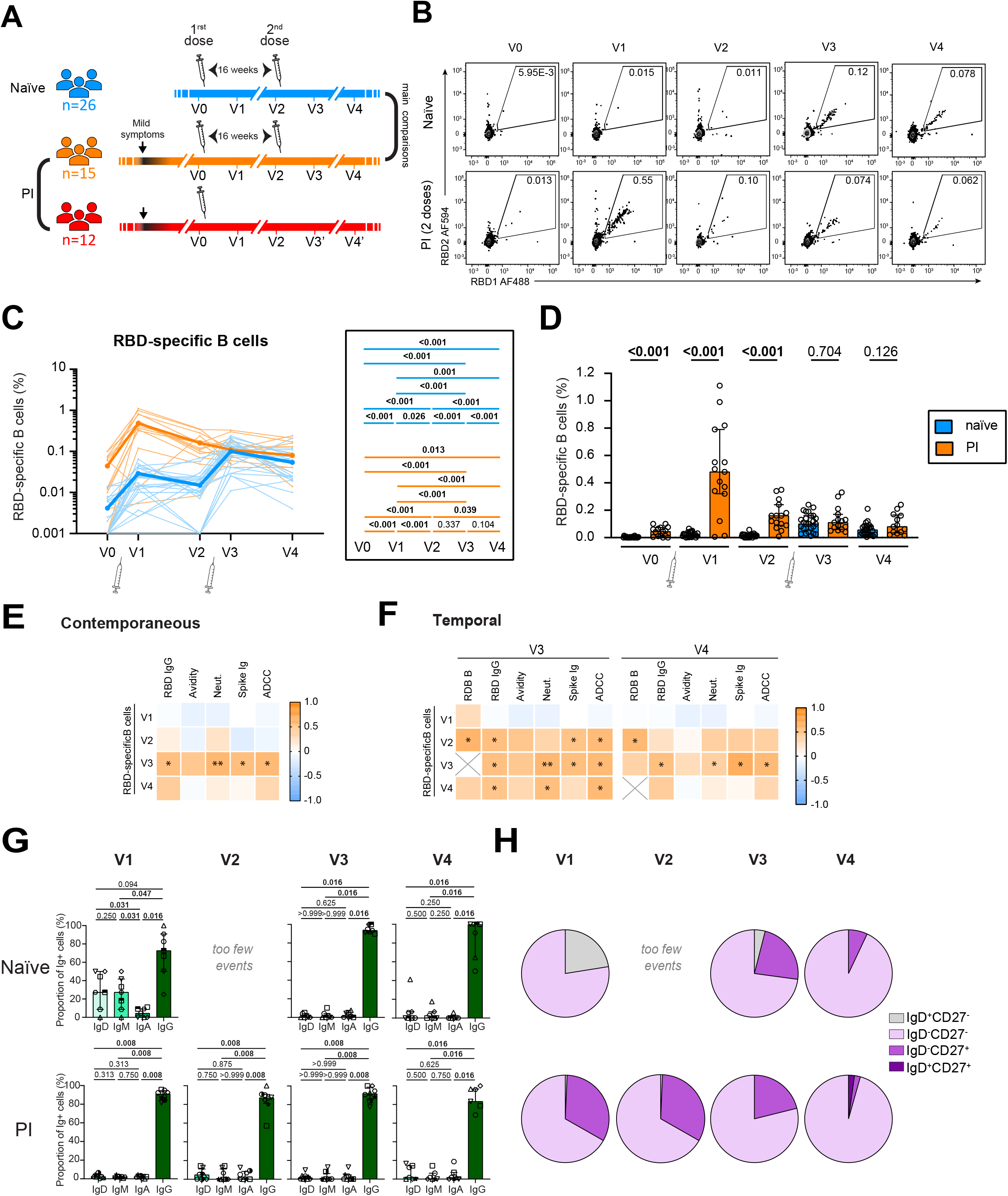
Marked differences in B cell responses to the first BNT162b2 dose between naïve and PI participants contrast with convergent features after boosting. **(A)** Schematic representation of study design, visits and vaccine dose administration (indicated by a syringe). Blood samples were collected at 5 time points: at baseline (V0); 3 weeks after the first dose (V1); 12 weeks after the first dose (V2); 3 weeks after the second dose for participants receiving two doses (V3) and 16 weeks after the second dose (V4). For participants receiving a single dose, V3’ was sampled 19 weeks after the first dose, and V4’ 16 weeks after the second dose. Actual times are summarized in Table 1. **(B)** Representative gating to identify RBD-specific B cells. **(CD)** Kinetics of RBD-specific B cell responses in previously naïve (blue) or pre-infected (PI; orange) participants receiving two vaccine doses. **(C)** Lines connect data from the same donor. The bold line represents the median value of each cohort. Right panel: Wilcoxon tests for each pairwise comparisons. **(D)** Comparisons between naïve and PI participants. The bars represent median and interquartile ranges. Mann-Whitney tests are shown. **(EF)** Heatmap showing **(E)** contemporaneous or **(F)** temporal correlations of RBD-specific B cells vs the indicated antibody responses (n=22). Significant correlations by Spearman tests are shown (*: *p* < 0.05, **: *p* < 0.01, ***: *p* < 0.001). **(G)** Frequencies of IgD, IgM, IgA and IgG-positive cells in RBD-specific memory B cells in naïve and PI donors, with Wilcoxon tests. **(H)** Proportion of IgD+/− and CD27+/− populations in RBD-specific memory B cells in naïve and PI donors. In GH, V2 for naïve participants could not be analyzed because of low number of events. In **CD**) n=26 naïve, n=15 PI; in **E**) n=26 naïve; in **GH**) n=7 naïve and n=8 PI.

**Table 1.** Clinical characteristics of the study participants ^†^.

| BNT162b2 vaccine | Previously naïve cohort | Previously-infected (PI) cohort |  |  |
| --- | --- | --- | --- | --- |
|  | Two doses* | Two doses* | Single dose ** | Entire (PI) cohort |
| Variable | (n=26) | (n=15) | (n=12) | (n=27) |
| Age | 51 (41-56) | 47 (43-56) | 51 (34-62) | 48 (39-59) |
| Sex |  |  |  |  |
| Male | 11 (42%) | 10 (66%) | 4 (33%) | 14 (52%) |
| Female | 15 (58%) | 5 (34%) | 8 (66%) | 13 (48%) |
| Previous SARS-CoV-2 infection |  |  |  |  |
| Days between days of symptom onset and first vaccine dose | NA | 274 (258-307) | 287 (227-306) | 281 (250-307) |
| Vaccine dose spacing |  |  |  |  |
| Days between doses 1 and 2 | 111 (109-112) | 110 (110-112) | NA | NA |
| Visits for immunological profiling |  |  |  |  |
| V0, days before first dose | 1 (0-5) | 24 (6-43) | 18 (7-45) | 23 (6-43) |
| V1, days after first dose | 21 (19-26) | 20 (19-21) | 20 (18-21) | 20 (18-21) |
| V2, days after first dose | 83 (82-84) | 89 (86-93) | 90 (87-94) | 89 (86-93) |
| V2, days before second dose | 28 (26-29) | 23 (18-28) | NA | NA |
| V3, days after first dose | 133 (130-139) | 138 (132-142) | 132 (130-138) | 136 (131-141) |
| V3, days after second dose | 21 (20-27) | 22 (18-28) | NA | NA |
| V4, days after first dose | 224 (222-228) | 224 (222-227) | 227 (223-237) | NA |
| V4, days after second dose | 112 (110-119) | 113 (110-117) | NA | NA |

<sup>†</sup> Values displayed are medians, with IQR: interquartile range in parentheses for continuous variables, or percentages for categorical variables.
\* The previously naïve cohort and previously-infected cohort that also received two vaccine doses were compared by the following statistical tests: for continuous variables, Mann-Whitney U test; for categorical variables, Fisher's test. Values highlighted in light grey are statistically different between the pre-infected naïve and pre-infected cohorts. No statistical difference was found between the two pre-infected subcohorts, except between naïve and pre-infected for days before V0 and days after V2.
\*\* The previously-infected cohort with 1 dose was likewise compared to the previously-infected cohort that received 2 doses. No significant differences was observed between the two cohorts.

### Marked differences in B cell responses to the first BNT162b2 dose between naïve and PI participants contrast with convergent features after boosting

To evaluate SARS-CoV-2-specific B cells, we focused on RBD to minimize inclusion of B cells cross-reactive to endemic coronaviruses (Hicks et al., 2021; Klumpp-Thomas et al., 2021). Co-detection of two fluorescently labeled recombinant RBD probes greatly enhances specificity (Figure 1B and (Anand et al., 2021); flow cytometry panel, Table S1; gating strategy, Figure S1A). We examined the magnitude of RBD-specific B cells (defined as RBD1^+^RBD2^+^CD19^+^CD20^+^) in the two-dose cohorts (Figure 1CD). In naïve individuals, we observed little baseline signal at V0 and the generation of a significant, albeit small, RBD-specific B cell population at V1. This primary response underwent partial attrition at the V2 memory time point. The second dose elicited a brisk recall response at V3. Responses subsequently declined at V4 yet remained significantly higher than at pre-boost timepoints. The pattern markedly differed in PI (Figure 1CD). Consistent with previous SARS-CoV-2 exposure, RBD-specific B cells were already present at V0. This response increased sharply at V1, followed by attrition at V2. We observed no boosting effect after the second dose, and no significant decline at V4. The response to the first BNT162b2 dose in PI (V1) differed from the second BNT162b2 dose in naïve (V3), as we observed differences in magnitude (Figure S1B). Therefore, the RBD-specific B cell kinetics between the two cohorts markedly differed after the first dose, converged after the second dose, and the subsequent moderate decline observed at V4 showed no significant difference (Figure 1D). In single-dose PI, we observed stable B cell responses at V3’ and V4’ compared to V2, consistent with a steady memory B cell pool after an initial decline between V1 and V2 (Figure S1C).

We next investigated the relationships between RBD-specific B cell frequencies at the different time points and antibody responses in naïve participants: RBD-specific IgG antibody levels, anti-RBD IgG avidity, neutralization activity, cell-binding ELISA (CBE) antibody levels, and antibody-dependent cellular cytotoxicity (ADCC) (Figure 1EF). RBD-specific B cells responses positively correlated with contemporaneous antibody responses at V3, but not at V1 (Figure 1E). Contemporary correlations were lost at V4. Early V1 B cell responses were not associated with subsequent V3 and V4 antibody responses, but significant correlations were found between RBD-specific B cells at V2 and RBD IgG, total Spike antibody, cell binding and ADCC at V3 (Figure 1F). Similarly, V3 RBD-specific B cell responses correlated significantly with V4 RBD-specific IgG, total anti-Spike antibody levels and ADCC, suggesting that the B cell pool post-boost conditioned the long-term quantity and quality of the humoral response.

To determine how B cell populations qualitatively evolved, we measured IgD, IgM, IgG and IgA expression profiles of RBD-specific B cells. In the naïve cohort, we detected subpopulations of IgD^+^, IgM^+^ and IgA^+^ cells at V1, whose proportion decreased at V3 and V4 visits. In contrast, RBD-specific memory B cells in PI donors were almost entirely IgG^+^ at all timepoints (Figure 1G, Figure S1EF). To assess B cell differentiation, we quantified IgD and CD27 co-expression (Figure S1G). CD27 is predominantly expressed on memory B cells (Tangye et al., 1998), and IgD on unswitched B cells (Moore et al., 1981). In the naïve cohort, IgD^+^CD27^-^ RBD-specific B cells present at V1 disappeared at V3, while IgD^-^CD27^+^RBD-specific B cells emerged (Figure 1H), consistent with isotype-switched memory B cells. This subset contracted at V4. In PI, IgD^-^CD27^+^ cells already present at baseline expanded after priming and remained stable at V2. Boosting did not further expand this subset. Instead, it gradually declined at V3 and V4. A class-switched IgG^+^ DN population dominated at all time points (Figure 1H, Figure S1HI).

These data show that despite the long 16-week interval and the divergent RBD-specific B cells trajectories after the first dose, boosting in naïve subjects induced robust recall responses with a mature phenotype that converged with those observed in PI individuals.

### The first and delayed second vaccine doses elicit Spike-specific CD4^+^ T cell responses of similar magnitude

CD4^+^ T cells help play a critical role in development of B cell and CD8^+^ T cell immunity. We measured Spike-specific T cell responses at the V0-V4 timepoints in the three cohorts (Figure 2 and S2). As in our previous work, we used a TCR-dependent activation induced marker (AIM) assay that broadly identifies antigen-specific T cells (Morou et al., 2019; Niessl et al., 2020a; Niessl et al., 2020b; Tauzin et al., 2021b) and functional profiling by intracellular cytokine staining (ICS) (Tauzin et al., 2021b)(flow cytometry panels: Tables S2-3).

**Figure 2.**
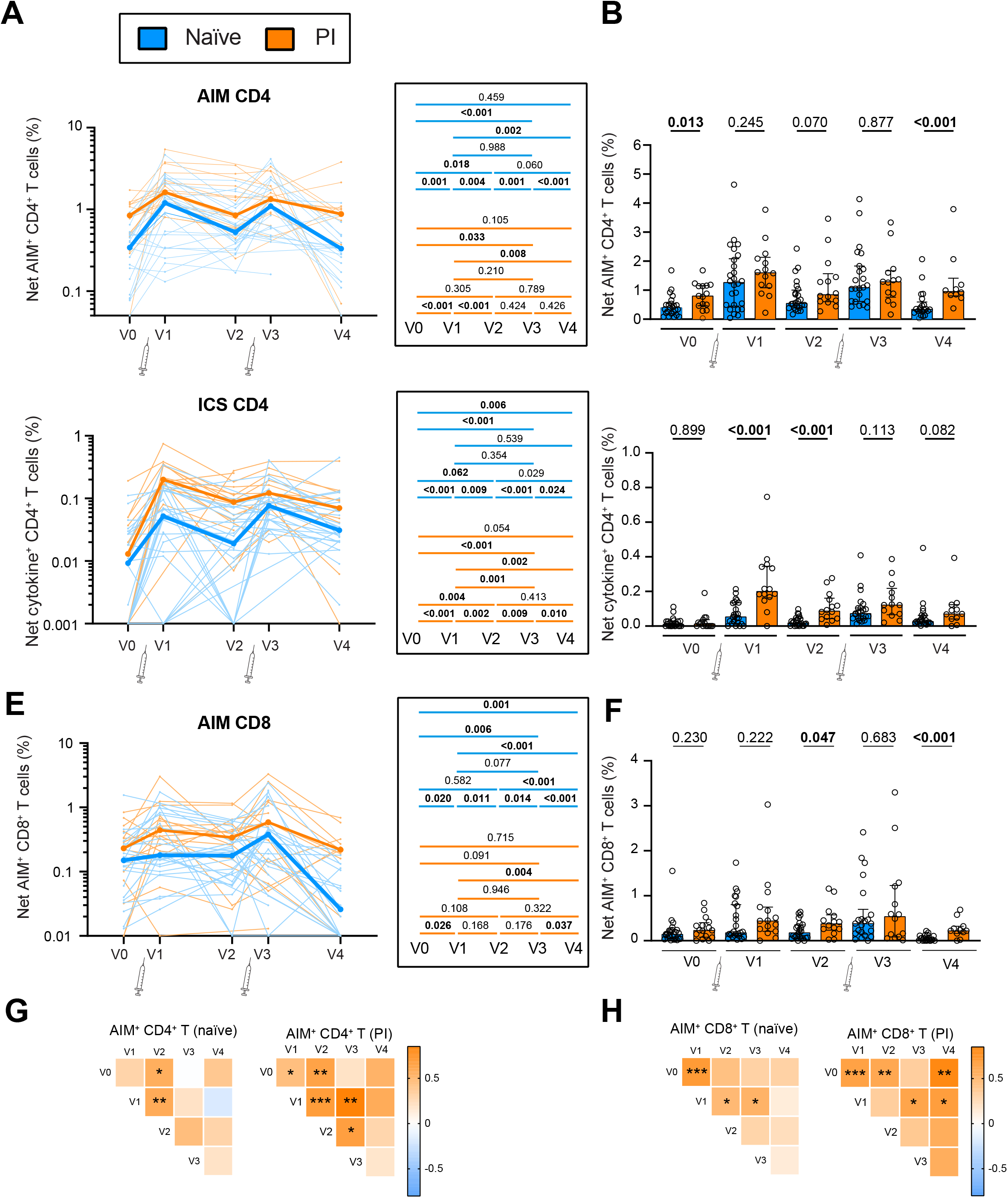
The first and delayed second vaccine doses elicit Spike-specific CD4^+^ T cell responses of similar magnitude. Frequencies of SARS-CoV-2 Spike specific CD4^+^ and CD8^+^ T cells in naïve (blue) and PI (orange) receiving two vaccine doses. PBMCs were stimulated *ex vivo* with a pool of overlapping Spike peptides. **(AB)** Net AIM^+^CD4^+^ T cell responses. **(A)** Longitudinal analysis of Spike-specific AIM^+^CD4^+^ T cell responses. Lines connect data from the same donor. The bold line represents the median value of each cohort. Right panel: Wilcoxon tests for each pairwise comparisons. **(B)** Cohort comparisons at each time point. The bars represent median and interquartile ranges. Mann-Whitney tests are shown. **(CD)** Evolution of the net cytokine^+^CD4^+^ T cell responses measured by ICS. **(C)** Longitudinal analysis of the magnitude of cytokine^+^CD4^+^ T cell responses. Lines connect data from the same donor. The bold line represents the median value of each cohort. Right panel: Wilcoxon tests for each pairwise comparison. **(D)** Cohort comparisons at each time point. The bars represent median and interquartile ranges. **(EF)** Net Spike-specific AIM^+^CD8^+^ T cell responses in naïve and PI participants. **(E)** Longitudinal AIM^+^CD8^+^ T cell responses. Lines connect data from the same donor. The bold line represents the median value of each cohort. Right panel: Wilcoxon tests for each pairwise comparison. **(F)**. Cohort comparisons at each time point. The bars represent median and interquartile ranges. In A, C and E, the syringe indicates vaccine dose inoculation. **(GH)** Heatmap showing temporal correlations of **(G)** AIM^+^CD4^+^ and **(H)** AIM^+^CD8^+^ T cells between the different time points for naïve and PI participants. Significant Spearman test results are indicated (*: *p* < 0.05, **: *p* < 0.01, ***: *p* < 0.001). In **A-F**) n=26 naïve, n=15 PI participants; in **GH**) n=26 naïve; n=27 PI (comparisons at time points V0, V1 and V2), n=15 PI (comparisons at time points V3 and V4).

The AIM assay involved a 15-h incubation of PBMCs with an overlapping peptide pool spanning the Spike coding sequence and the upregulation of CD69, CD40L, 4-1BB and OX-40 upon stimulation. We used an AND/OR Boolean combination gating to assess total frequencies of antigen-specific CD4^+^ and CD8^+^ T cells (Figure S2AB) (Niessl et al., 2020a; Tauzin et al., 2021b). At V3, all individuals had CD4^+^ T cell responses (Figure S2C), and most had CD8^+^ T cell responses (Figure S2D).

In contrast to B cell responses, the kinetics of Spike-specific AIM^+^CD4^+^ T cell responses were more similar between naïve and PI individuals (Figure 2A). Several naïve participants had detectable AIM^+^CD4^+^ T cell responses at baseline, probably due to cross-reactivity with other coronaviruses (Mateus et al., 2020). The significant increase at V1 was followed by a moderate attrition at the V2 memory timepoint. The second dose significantly boosted the responses at V3 in naïve, whereas the increase was non-significant in PI. The significant difference in median magnitude of AIM^+^CD4^+^ T cell responses between naïve and PI became non-significant at V3 before increasing again at V4 due to a faster decay of these Thelper responses in naïve compared to PI (Figure 2B).

The ICS assay involved a 6-h stimulation with the Spike peptide pool and measurement of the cytokines and effector molecules IFN-γ, IL-2, TNF-α, IL-17A, IL-10 and CD107a. We determined total cytokine+ CD4^+^ T cell responses by an AND/OR Boolean combination gating strategy (Figure S2E). The ICS patterns in both cohorts largely paralleled the AIM assays, albeit at a lower magnitude (Figure 2CD and S2F). Consistent with the lower ability of ICS to detect memory cells compared to recently primed or reactivated cells (da Silva Antunes et al., 2018), the relative increase in cytokine^+^CD4^+^ T cells was stronger at V1 vs V0 and V3 vs V2. Cytokine^+^CD4^+^ T cell responses at V4 remained significantly higher than at baseline, showing longer-term memory, but without significant gain compared to V2.

The magnitude of Spike-specific AIM^+^ CD8^+^ T cell responses was significantly lower than that of Spike-specific AIM^+^CD4^+^ T cell responses at all time points (Figure 2EF and, S2G). In naïve, but not in PI, the second vaccine dose significantly increased AIM^+^CD8^+^ T cell responses (Figure 2E). AIM^+^CD8^+^ T cell responses declined significantly at V4 for both cohorts. Total cytokine^+^CD8^+^ T cell responses were weak or undetectable in most participants at all time points, precluding their detailed analysis (Figure S2H).

To define the evolution of T cell responses in the absence of boosting, we examined the single-dose PI cohort. The magnitude of CD4^+^ (Figure S2I) and CD8^+^ T cell (Figure S2J) responses did not further decline at V3, suggesting stable memory.

As expansion of previously primed antigen-specific T cells may impact T cell responses to vaccination, we examined correlations across visits (Figure 2G) and found a significant association or a strong trend between CD4^+^ T cell responses at V0 and the post-first dose time points V1 and V2, but not after the second dose. Similar to CD4^+^ T cells, we observed that preexisting CD8^+^ T cell responses at V0 significantly correlated to responses to the first dose at V1, and that this association disappeared after the second dose (Figure 2H). Therefore, the second vaccine dose reduced the heterogeneity in magnitude of T cell responses and their links to pre-vaccination immunity.

These data show that a single dose of the BNT162b2 is sufficient to induce CD4 Thelper responses, and CD8^+^ T cell responses in most participants. After a 16-week interval, the second dose boosts CD4^+^ and CD8^+^ T cell responses back to the peak magnitudes reached soon after priming. Pre-vaccination T cell immunity is associated with the BNT162b2 -induced CD4^+^ and CD8^+^ T cell responses to the first vaccination, but this correlation is lost after boosting.

### The 16-week interval BNT162b2 regimen elicits phenotypically diverse CD4^+^ T helper subsets

We next profiled the qualitative heterogeneity and evolution of Spike-specific AIM^+^CD4^+^T cells. To avoid *a priori* defined marker combinations, we performed unsupervised analyses of the high-dimensional flow cytometric phenotyping data (Figure 3). We examined chemokine receptors that are preferentially, but not exclusively, expressed by some lineages and involved in tissue homing (CXCR5 for Tfh; CXCR3 for Th1; CCR6 for Th17/Th22 and mucosal homing; CXCR6 for pulmonary mucosal homing (Day et al., 2009; Morgan et al., 2015)), CD38 and HLA-DR as activation markers, and PD-1 as inhibitory checkpoint.

**Figure 3:**
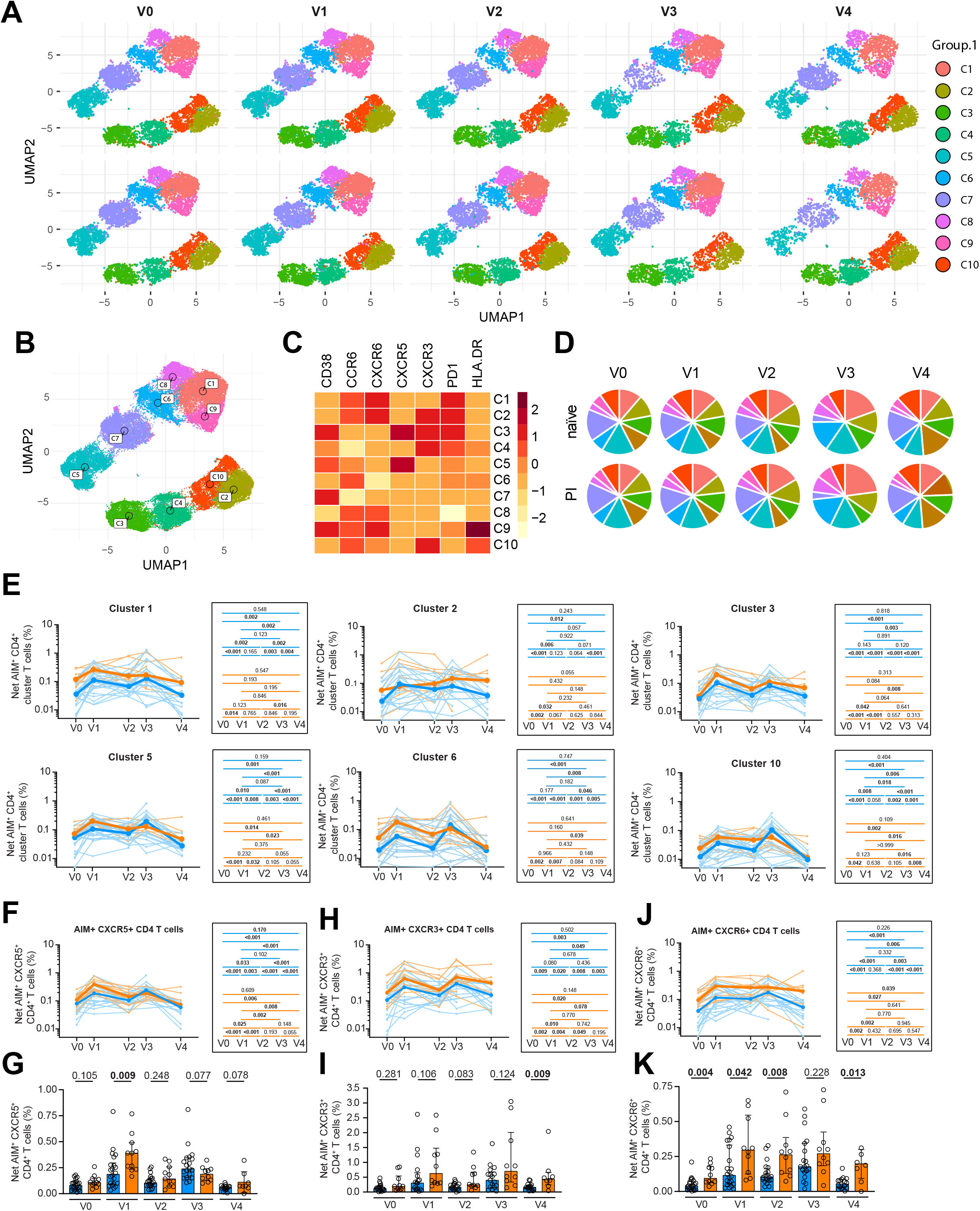
The 16-week interval BNT162b2 regimen elicits phenotypically diverse CD4^+^ T helper subsets. **(A)** Multiparametric UMAP representation of Spike-specific AIM^+^CD4^+^ T cells based on CXCR3, CXCR5, CXCR6, CCR6, PD1, HLA-DR and CD38 expression at each time point, aggregated data for the two-dose naïve and PI cohorts. The colors identify 10 populations clustered by unsupervised analysis using Phenograph. **(B)** Each cluster are labeled on the global UMAP. **(C)** Heat map summarizing for each cluster the mean fluorescence intensity of each loaded parameter. **(D)** Pie charts depicting the representation of each identified cluster within total AIM^+^CD4^+^ T cells. **(E, F,H,J**) Longitudinal frequencies of selected AIM^+^CD4^+^ T cell **(E)** clusters and **(F)** CXCR5-, **(G)** CXCR6-, **(H)** CXCR3-expressing AIM^+^ CD4^+^ T cells. Lines connect data from the same donor. The bold line represents the median value of each cohort. Right panel: Wilcoxon tests for each pairwise comparison. **(G,I,K)** Cohort comparisons at each time point for **(G)** CXCR5-, **(I)** CXCR3 -, **(K)** CXCR6-expressing AIM^+^ CD4^+^ T cells. The bars represent median and interquartile ranges. In A-K) n=22 naïve, n=11 PI participants.

We illustrated the distribution of clustered populations by the uniform manifold approximation and projection (UMAP) algorithm (Becht et al., 2018). Cluster identity was performed using Phenograph (Levine et al., 2015), resulting in the identification of 10 clusters (Figure 3AB) based on distinct profiles of relative marker expression (Figure 3C, and S3A). All 10 clusters were detectable at V0 and persisted at all timepoints. The relative frequencies of each cluster did not show major differences across visits (Figure 3D), but there were fluctuations and interindividual variations within cohorts (Figure 3E, Figure S3B). We did not observe emergence of new Thelper clusters after the second inoculation. While variability and relatively small cohort size precluded definitive conclusions about the behavior of individual clusters, some general trends were observed. In naïve, most clusters showed either a significant or a trend for increase after the second dose (Figure 3E), except C4 (Figure S3B). These included clusters enriched in CXCR5 (C3 and C5) and CXCR3 (C2, C3 and C10). In contrast, the qualitative response to the second in PI was more constrained (Figure 3D and S4C). Consistent with the analysis on total AIM^+^ T CD4^+^ cells, all naïve participant clusters declined at V4, excepted C4 (Figure 3E and S3B). Although some clusters also showed trend for decline in PI (C5, C6, C9 and C10), most did not (C1, C2, C3, C4, C7 and C8).

We next performed univariate analyses of chemokine receptor expression (Figure 3F-K and S3D-I). CXCR5^+^AIM^+^CD4^+^ T cell did not further expand after the second compared to first dose, and trajectories did not statistically differ between cohorts past V1. CXCR3^+^AIM^+^CD4^+^ T cell increased similarly after either dose, with significant decline post first inoculation. This pattern was similar in naïve and PI, but the CXCR3^+^ subset was more abundant_IgD_ in PI. CXCR6^+^AIM^+^ and CCR6^+^AIM^+^CD4^+^ T cells remained persistently elevated after priming in PI, while in naïve they were weaker at early timepoints but responsive to the second dose at V3. However, they declined at the late memory timepoint V4 in this cohort.

Therefore, the first vaccine dose already elicits phenotypically diverse Thelper clusters that do not necessarily fit with canonical lineages. The second vaccine dose variably impacted these subsets but did not elicit new clusters. Thelper phenotype in PI was enriched in markers suggestive of prior mucosal priming.

### The delayed second BNT162b2 dose leads to partially convergent functional profiles in naïve and PI participants

We next applied the same unsupervised analysis pipeline to Spike-specific cytokine^+^CD4^+^ T cells for the six functions measured, identifying 11 functional clusters (Figure 4ABC and S4A) that were present at all timepoints (Figure 4AD), with notable inter-individual differences within each cohort (Figure 4E and S4B). However, we observed clearer functional differences between cohorts and between doses when compared to phenotypic analysis. Most clusters increased after both doses in naïve, whereas they expanded only after the first dose in PI (Figure 4E and S4B). Individual clusters followed different trajectories, depending on preinfection status (Figure 4E, and S4C). The evolution of the IL-2 enriched C1, the most abundant cluster, was similar in the PI and naïve cohorts, except for significant contraction at V4 in naïve only. In contrast, the C2 and C3 clusters, characterized by high IFN-γ expression, were markedly larger in PI after the first dose, but responded more to the second dose in naïve. Consequently, the responses of C2 and C3 partially converged at V3 compared to V1; they significantly contracted at V4. The polyfunctional cluster C5, enriched in IFN-γ, IL-2, TNF-α and CD107a, showed yet another pattern: it was expanded in PI compared to naïve at all timepoints, and showed excellent long-term stability in both cohorts.

**Figure 4:**
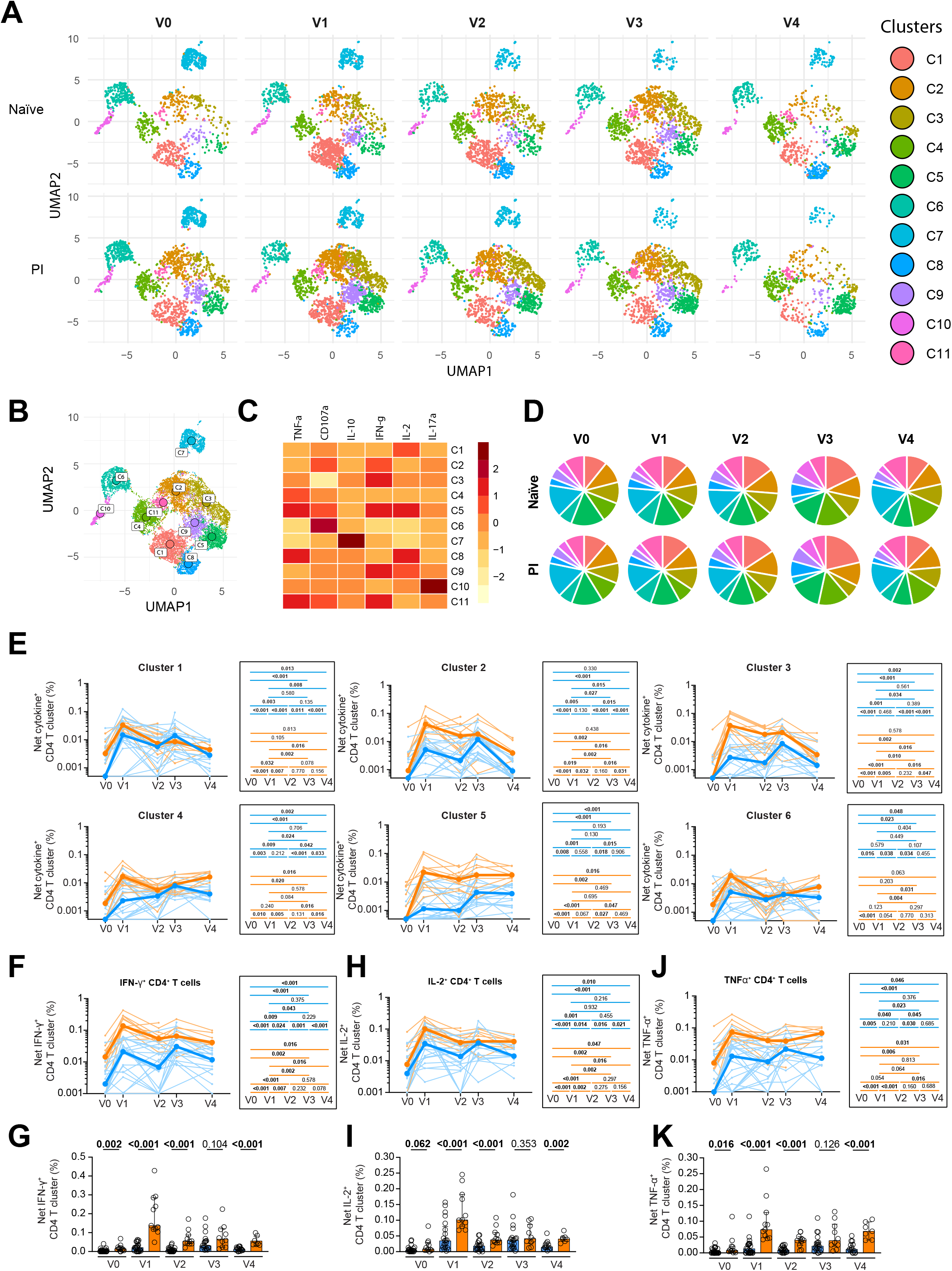
The delayed second BNT162b2 dose leads to partially convergent functional profiles in naïve and PI participants. **(A)** Multiparametric UMAP representation of Spikespecific ICS cytokine^+^CD4^+^ T cells based on IFN-γ, IL-2, TNF-α, IL-17A, IL-10 and CD107a expression at each time point, aggregated data for the two-dose naïve and PI cohorts. The colors identify 11 populations clustered by unsupervised analysis using Phenograph. **(B)** Each cluster is labeled on the global UMAP. **(C)** Heat map summarizing for each cluster the mean fluorescence intensity of each loaded parameter. **(D)** Pie charts depicting the representation of each identified cluster within total cytokine^+^CD4^+^ T cells. **(E, F, H, J**) Longitudinal frequencies of selected cytokine^+^CD4^+^ T cell **(E)** clusters and **(F)** IFN-γ^+^, **(H)** IL-2^+^, **(J)** TNF-α^+^ single functions in naïve (blue) and PI (orange) participants. Lines connect data from the same donor. The bold line represents the median value of each cohort. Right panel: Wilcoxon tests for each pairwise comparison. **(G, I, K)** Cohort comparisons at each time point for **(G)** IFN-γ^+^, **(I)** IL-2^+^, **(K)** TNF-α^+^ single functions. The bars represent median and interquartile ranges. In A-K) n=22 naïve, n=11 PI participants.

Single-parameter analyses (Figure 4F-K and S4DE) showed that the first dose significantly increased IFN-γ^+^, IL-2^+^ and TNF-α^+^ CD4+ T cell responses, particularly for PI. The second dose significantly boosted these responses in naïve only, contrasting with little effect in PI. These differential trajectories led to partially convergent CD4^+^ T cell functions after repeated antigenic challenges in naïve and PI at V3. However, consistent with AIM measurements, weaker responses in naïve at V4 led to re-emergence of significant differences at this late memory timepoint.

These analyses show that pre-infection status is associated with significant differences in the functional profile elicited by the first BNT162b2 dose. Preferential expansion of Th1-cytokine-enriched subsets after boosting in naïve participants contrasting with stable responses in PI leads to partial, and possibly transient, convergence of Thelper functions between cohorts after full vaccination. The unsupervised analysis reveals a polyfunctional cluster of CD4^+^ T cell stably maintained at the late memory timepoint.

### Temporal relationships between antigen-specific CD4^+^ T cell, B cell and CD8^+^ T cell responses

As CD4^+^ T cell help is essential for optimal adaptive B cell and CD8^+^ T cell immunity, we next examined the temporal associations between these immune components (Figure 5). We considered 23 CD4^+^ T cell subpopulations at the V0-V3 time points: the total Spike-specific AIM^+^CD4^+^ T cells, the AIM^+^C1-C10 clusters, the total Spike-specific cytokine^+^CD4^+^ T cells and the ICS (cyto^+^C1-C11) clusters. We applied unsupervised clustering analyses to determine the longitudinal relationships between these Thelper subsets and RBD-specific B cell (Figure 5A) and AIM^+^CD8 T cell (Figure 5B) responses, measured at V3 after completion of the vaccination regimen.

**Figure 5:**
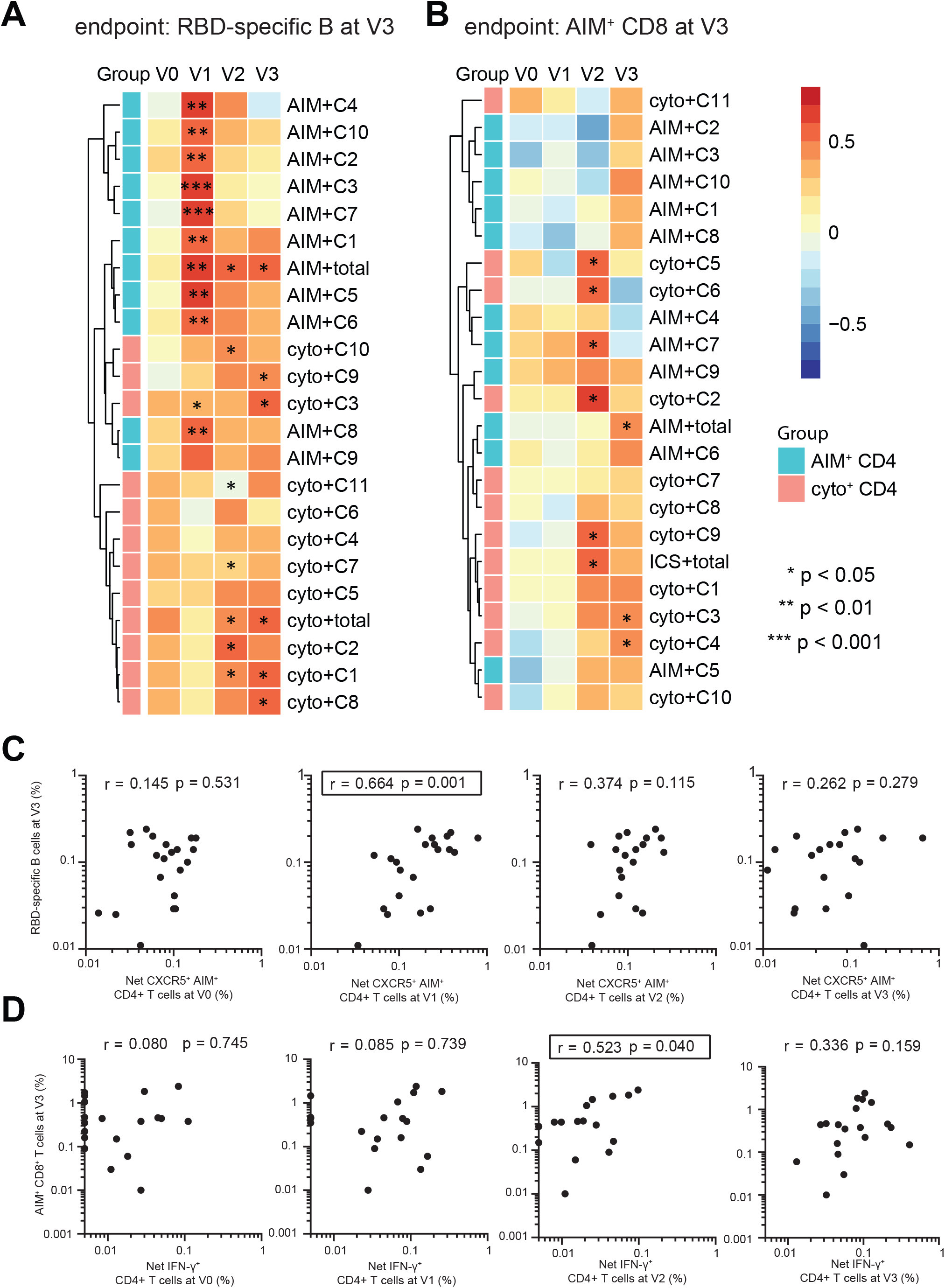
Temporal relationships between antigen-specific CD4^+^ T cell, B cell and CD8^+^T cell responses in naïve participants. **(AB)** Heat maps displaying temporal correlations between the different subsets of Spike-specific CD4^+^ T cells measured by AIM or ICS assays at the V0, V1, V2 and V3 timepoints and **(A)** RBD-specific B cell frequencies measured at V3; **(B)** AIM^+^CD8^+^ T cell frequencies measured at V3. Asterisks indicate the significance (*: *p* < 0.05, **: *p* < 0.01, ***: *p* < 0.001). The CD4^+^ T cell clusters were stratified by assay (cyan = AIM, light red= ICS). (**C**) Correlations between frequencies of AIM^+^CXCR5^+^ CD4^+^ T cells (for cTfh) at the V0-V3 visits and RBD-specific B cell frequencies at V3. (**D**) Correlations between frequencies of IFN-γ+ (as Th1 function) at the V0-V3 visits and AIM^+^CD8^+^ T cell at V3. The r and p values from a Spearman test are indicated in each graph. **A,D**) n=21 naïve participants; **B,D**) n=19.

We observed significant positive correlations between all AIM^+^CD4^+^ T cell subsets elicited at V1 and the B cell responses at V3 after the second dose (Figure 5A). This contrasted with the weaker correlations between V2-V3 cytokine+ Thelper responses and RBD-specific B cells at V3. Some of the positively correlated clusters (AIM^+^C3, AIM^+^C5) were enriched in CXCR5^+^ cells. Consistently, CXCR5^+^AIM^+^CD4^+^ T cell responses at V1 strongly correlated with B cell responses at V3, but this association weakened for V2 and disappeared at V3 (Figure 5C). Similar patterns were seen with total AIM^+^CD4^+^ T cells (Figure S5A) and for some non-cTfh subsets, but we did not have the statistical power to rank the strength of the correlations.

We next examined the temporal associations between longitudinal Thelper subsets and AIM^+^CD8^+^ T cells at V3 (Figure 5B). Thelper responses at V1 showed no significant correlation with AIM^+^CD8^+^ T cells at V3. However, we found significant correlations between cytokine^+^CD4^+^ T cell subsets at the pre-boost V2 memory timepoint or at the contemporaneous V3 and the AIM^+^CD8^+^ T cell responses at V3, and IFN-γ^+^CD4^+^ T cells at V2 correlated with AIM^+^CD8^+^ T cells at V3 (Figure 5D), as did total cytokine^+^CD4^+^ T cell responses (Figure S5B).

The differential temporal associations between antigen-specific CD4^+^ T cell, B and CD8^+^ T cell immunity, suggest different requirements for the coordination of these responses.

### Immune profile kinetics in naïve and PI vaccinees shows only partial, and transient, convergence after the delayed second dose

Our data suggest that the relationships between the different immune parameters after the first vaccine dose were strongly influenced by prior infection history, while its impact decreased, but did not disappear, after the second dose. We therefore performed an integrated analysis of 34 features of antibody, B cell CD4^+^ T cell, CD8^+^ T cell responses (Figure 6A). V1 to V4 time points were first loaded altogether (Figure 6B), then from this master PCA we depicted each timepoint separately (Figure 6C). The two cohorts clustered apart at V1 due to a significant difference in PC1 (Figure 6D). The distance between groups decreased upon attrition of the responses (V2). No statistical difference between naïve and PI PC1 was observed at V3, showing convergence of the immune features. Importantly, however, the PC1-driven distinction between naïve and PI reemerged at the late memory time point V4.

**Figure 6:**
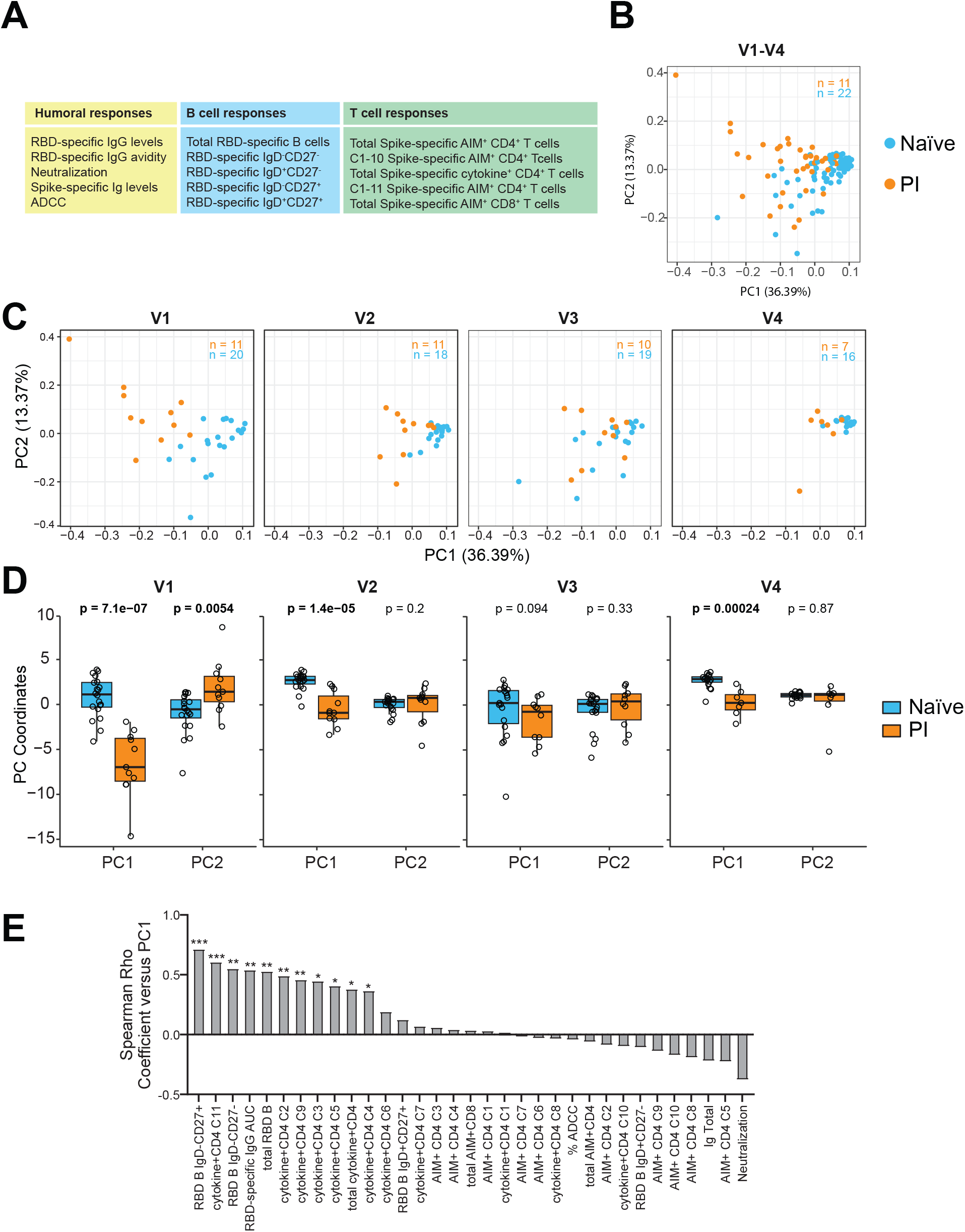
Immune profile kinetics between naïve and PI vaccinees shows only partial, and transient, convergence after the delayed second dose. Integrated PCA analysis combining various immune features to compare evolution of vaccine responses in the two-dose naïve and PI cohorts. **(A)** List of the 34 antigen-specific immune magnitudes included in the PCA analysis. (**B)** Global PCA analysis. The percentage on the x and y axes present the variance attributed to PC1 and PC2, respectively. **(C)** PCA sub-analyses divided by time points. Only complete datasets could be loaded in the PCA analysis. The numbers of participants by PCA are shown in each plot. **(D)** Box and whisker plots of the PC1 and PC2 between group are with Mann-Whitney tests. In **BC**, blue is representing naïve participants and orange, PI. **(E)** Spearman correlations between each individual immune feature and PC1 (* *p* < 0.05; ** *p* < 0.01; *** *p* < 0.001).

We sought to identify the features underlying the group clustering at V1 using the same approach focused on AIM^+^CD4^+^ and cytokine^+^CD4^+^ T cell responses. The correlation between the immune features and PC1 identified anti-RBD IgG levels, memory RBD-specific B cells, IFN-γ-enriched Spike-specific CD4^+^ T cell clusters with little contribution of AIM^+^ CD4 T cells. A PCA analysis performed using AIM^+^ CD4^+^ T cell features confirmed the limited contribution of these features to cohort clustering (Figure S6A), in contrast to the cytokine+ CD4^+^ T cells (Figure S6B).

Therefore, unsupervised integrated analysis shows that pre-infection status shapes vaccine-induced immunity after the first dose, while its influence largely wanes in the shortterm response to the second dose but subsequently becomes more manifest again 8 months after initial inoculation.

## DISCUSSION

The decision to extend intervals between doses of the BNT162b2 mRNA vaccine led to concerns about vaccine immunogenicity and efficacy. Here, we profiled the B cell, CD4^+^ T cell, CD8^+^ T cell and antibody responses in SARS-CoV-2 naïve and PI individuals who received the two vaccine doses 16 weeks apart. We longitudinally followed these immune features from baseline over an 8-month period to determine the characteristics and temporal associations of the immune features elicited by this wide-interval immunization regimen.

We observed that in naïve participants, the priming dose elicited RBD-specific responses of low magnitude at three weeks, which slightly declined at a 12-week memory time point. However, administration of the second dose after the 16-week interval strongly enhanced these B cell responses, which after having acquired phenotypic features of isotype-switched memory B cells underwent only a moderate contraction in the following months. These robust B cell responses were associated with the development of strong and broad humoral responses, as we reported (Tauzin et al., 2021a). The phenotypic changes were consistent with B cell maturation: immunoglobulin isotype expression was dominated by IgG^+^at all time points, and included IgD^+^, IgM^+^ and IgA^+^ cells after the first dose that were mostly absent after the second dose; and the differentiation phenotype was characterized by expansion of a CD27^+^ IgD^-^ memory subset after the second dose compared to the first dose. At all time points, a majority of double negative (CD27^-^ IgD^-^) cells was observed. Other studies have described this phenotypic subset in autoimmune diseases (Jenks et al., 2018; Wei et al., 2007) and in response to vaccination (Ruschil et al., 2020). Their transcriptional program is distinct from canonical switched memory cells and naïve cells (Jenks et al., 2018). They may be associated with an extra-follicular maturation pathway (Ruschil et al., 2020). In our study, the long-lasting persistence of these cells after boosting and their expression of RBD-specific IgG may suggest an atypical switched memory subset. These data are consistent with development of functional memory B cells with robust recall potential, alleviate the concern that an extended-interval regimen would lead to poor antibody immunity, and are in line with recent findings (Parry et al., 2021; Payne et al., 2021).

The kinetics of RBD-specific B cell responses differed in PI compared to naïve individuals. The first vaccine dose elicited a quantitatively small response in naïve but a brisk expansion in PI, with subsequent partial attrition before the delayed boost. Post-boost, B cell responses were similar in naïve and PI individuals following a strong expansion in naïve, but unchanged levels in PI. The responses after the first dose are in agreement with other studies of cellular and humoral responses (Efrati et al., 2021; Stamatatos et al., 2021; Tauzin et al., 2021b; Urbanowicz et al., 2021). A previous study showed a profound impact of the second dose on antigen-specific B cell responses in naïve participants, but a limited one in PI with the standard short-interval regimen, with convergent trajectories between the two groups (Goel et al., 2021). Our results demonstrate that this holds true even after an extended 16-week interval, consistent with the limited quantitative and qualitative enhancement of humoral immunity we documented in another report (Tauzin et al., 2021a).

The CD4^+^ T cell responses generated by vaccination were already quantitatively robust and phenotypically and functionally diversified after the first dose. Although these CD4^+^ T cell responses varied in magnitude among donors, they were reliably induced in all individuals examined. Mirroring the B cell findings and in contrast to the standard 3-week interval schedule, CD4^+^ T cell responses decrease moderately and transition to a memory phenotype before boosting. While the second dose significantly increased Spike-specific CD4^+^ T cells, these responses did not surpass those elicited by the first dose.

Unsupervised and supervised analyses demonstrated that the BNT162b2 vaccine elicits a highly diversified CD4^+^ T cell response, which is maintained over time with no novel distinct subset identified by our approach after the boost. Some qualitative Thelper features however, differed between the naïve and PI cohorts for the same point, and evolved between timepoints within cohorts. We observed higher frequencies of CCR6^+^ and CXCR6^+^ Thelper cells in PI than in naïve. As CXCR6, the CXCL16 ligand, is a homing molecule to the respiratory mucosa (Day et al., 2009; Morgan et al., 2015), these results are consistent with prior priming of CD4^+^ T cells at this anatomic site during SARS-CoV-2 infection in PI participants resulting in differences in their differentiation program, compared to intramuscular vaccine injection. We observed a similar pattern for CCR6, a marker of Th17 and Th22 cells that play an important role in maintaining mucosal barriers and contribute to pathogen clearance at mucosal surfaces (Aujla et al., 2008; Khader et al., 2007). Functional CD4^+^ T cell subsets also presented differential kinetics between naïve and PI individuals. While these differences between cohorts were stronger after the first dose, particularly for the Th1/cytotoxic clusters that were robust in PI, there were persistent late differences at the 8-month (V4) timepoint. In both cohorts, we identified a polyfunctional Th1 cell subset with excellent temporal stability. An analogous population has been associated with vaccine protection in a murine Leishmania model (Darrah et al., 2007).

Intriguingly, vaccination in PI, which corresponds immunologically to a heterologous prime-boost regimen, resulted in less decline after peak and significantly stronger CD4^+^ and CD8^+^ T cell responses at the late memory point than in naïve, which may suggest more durable immune memory. The single-dose PI group also exhibited excellent persistence of T cell memory up to 8 months after the single injection. The pattern was particularly pronounced for AIM^+^CD8^+^ T cells. Besides actual loss of CTL responses, there are other possible explanations: the measurements in peripheral blood may not reflect persistent tissue-resident memory populations in other anatomic compartments; and the activation-induced markers used may be unsensitive to identify some antigen-specific T cell subsets.

Subsets enriched in Tfh markers were already robustly induced early after priming and present at all timepoints, suggesting excellent induction by the BNT162b2 vaccine of this lineage critical for B cell help. Consistent with this notion, we identified strong temporal associations between several subsets of early vaccine-induced cTfh (and other CD4^+^ T cell subsets) after the first dose and B cell responses measured several months later, after the boost. This correlation was lost when CD4^+^ T cell responses were examined at later time points. While the observed disconnect in peripheral blood measurements at the late time points might be related to compartmentalization in lymphoid tissue (rather than major changes in CD4-B cell interplay), they suggest that the early antigen-specific CD4^+^ T cell responses critically shape the B cell pool, which will later respond to the delayed boosting. Despite the difference in dosing intervals, these results are thus overall consistent with the immune dynamics observed in the standard regimen (Oberhardt et al., 2021; Painter et al., 2021).

In contrast, Thelper responses with Th1 features identified at the early memory time point were better associated with the CTL responses after full vaccination, and we noted contemporaneous correlations as well. While this might suggest that the responsiveness of CD8^+^ T cells to boosting benefit from the pre-existing memory Thelper pool, mechanistic studies in murine models have shown that CD4^+^ T cell help is key at the time of CTL priming (Laidlaw et al., 2016), although they still play important roles later (Nakanishi et al., 2009). Our observational study does not allow to delineate causation from correlations due to other factors.

While the initial rationale of delaying the second dose was to provide some level of immunity more rapidly to a larger number of people in the context of limiting vaccine supply, our results suggest that this strategy could have the additional benefit of providing strong, multifaceted B and T cell immunity. The potential immunological benefits of increasing the interval between doses must be weighed against a prolonged period of good but still suboptimal protection, particularly while the virus and its different variants of concern are still circulating in the population at epidemic levels. Furthermore, these observations made in a HCW cohort may not be generalizable to vulnerable groups, particularly immunocompromised or elderly populations, in which the immune responses and the risk/benefit ratio may differ. Many countries now recommend a third dose, usually at least 6 months after the second dose. The benefit of a third dose in the context of a 16-week interval between the first and second dose will warrant further investigation.

## ACKNOWLEDGMENTS

The authors are grateful to the study participants. We thank the CRCHUM BSL3 and Flow Cytometry Platforms for technical assistance. We thank Dr. Stefan Pöhlmann (Georg-August University, Germany) for the plasmid coding for SARS-CoV-2 S glycoproteins and Dr. M. Gordon Joyce (U.S. MHRP) for the monoclonal antibody CR3022. This work was supported by a FRQS Merit Research Scholar award to D.E.K, the Fondation du CHUM, le Ministère de l’Économie et de l’Innovation du Québec, Programme de soutien aux organismes de recherche et d’innovation (to A.F). This work was also supported by CIHR operating grant # 178344 (D.E.K and A.F), foundation grant #352417 (A.F), by a CIHR operating Pandemic and Health Emergencies Research grant #177958 (A.F), an Exceptional Fund COVID-19 from the Canada Foundation for Innovation (CFI) #41027 to A.F. and D.E.K. Support was also provided by NIH AI144462 CHAVD to D.E.K. (Consortium PI: Dennis Burton). A.F. is the recipient of Canada Research Chair on Retroviral Entry no. RCHS0235 950-232424. V.M.L. is supported by a FRQS Junior 1 salary award. G.S is supported by scholarship from the Department of Microbiology, Infectious Disease and Immunology of the University of Montreal. G.B.B. was supported by a FRQS doctoral fellowship, R.G. and A.L. by a MITACS Accélération postdoctoral fellowship and J.P., by a CIHR doctoral fellowship. The funders had no role in study design, data collection and analysis, decision to publish, or preparation of the manuscript. We declare no competing interests.

## AUTHOR CONTRIBUTIONS

M.N., M.D, A.F and D.E.K. designed the studies. M.N, G.S, A.N, N.B and M.L. performed B cell and T cell assays. M.N, M.D. G.S, A.N, and O.T. performed and analyzed the B cell and T cell experiments. J.N. contributed to the T cell assay desing. O.T. performed unsupervised clustering analyses. L.M, A.T., G.B.B, D.V, S.Y.G, M.B, R.G, J.P., C.B., G.G.L. and J.R performed ELISA, ADCC, flow cytometry, avidity and neutralization assays. A.L., C.B., G.G.L., H.M., L.G., C.M., P.A., R.R., G.C.M., C.T. and V.M.L. secured and processed blood samples. G.G. produced and purified proteins. M.N., M.D., and D.E.K. wrote the manuscript with inputs from others. Every author has read, edited and approved the final manuscript.

## STAR METHODS

### RESOURCE AVAILABILITY

#### Lead contact

Further information and requests for resources and reagents should be directed to and will be fulfilled by the lead contact, Daniel E. Kaufmann.

#### Materials availability

All unique reagents generated during this study are available from the Lead contact upon a material transfer agreement (MTA).

#### Data and code availability

The published article includes all datasets generated and analyzed for this study. Further information and requests for resources and reagents should be directed to and will be fulfilled by the Lead Contact Author. The codes will be available as supplemental material.

### EXPERIMENTAL MODEL AND SUBJECT DETAILS

#### Ethics Statement

All work was conducted in accordance with the Declaration of Helsinki in terms of informed consent and approval by an appropriate institutional board. Blood samples were obtained from donors who consented to participate in this research project at CHUM (19.381).

#### Participants

No specific criteria such as number of patients (sample size), clinical or demographic were used for inclusion, beyond PCR confirmed SARS-CoV-2 infection in adults enrolled in the previously infected cohorts. Clinical data is summarized in Table 1.

#### PBMCs and plasma collection

PBMCs were isolated from blood samples by Ficoll density gradient centrifugation and cryopreserved in liquid nitrogen until use. Plasma was collected, heat-inactivated for 1 hour at 56°C and stored at −80°C until ready to use in subsequent experiments. Plasma from uninfected donors collected before the pandemic were used as negative controls and used to calculate the seropositivity threshold in our ELISA and ADCC assays.

#### Cell lines

293T human embryonic kidney and HOS cells (obtained from ATCC) were maintained at 37°C under 5% CO_2_ in Dulbecco’s modified Eagle’s medium (DMEM) (Wisent) containing 5% fetal bovine serum (FBS) (VWR) and 100 μg/mL of penicillin-streptomycin (Wisent). CEM.NKr CCR5+ cells (NIH AIDS reagent program) were maintained at 37°C under 5% CO_2_ in Roswell Park Memorial Institute (RPMI) 1640 medium (Gibco) containing 10% FBS and 100 μg/mL of penicillin-streptomycin. 293T-ACE2 cell line was previously reported (Prevost et al., 2020). HOS and CEM.NKr CCR5+ cells stably expressing the SARS-CoV-2 S glycoproteins (CEM.NKr.Spike cells) were previously reported (Anand et al., 2021).

### METHOD DETAILS

#### Protein expression and purification

FreeStyle 293F cells (Thermo Fisher Scientific) were grown in FreeStyle 293F medium (Thermo Fisher Scientific) to a density of 1 x 10^6^ cells/mL at 37°C with 8 % CO_2_ with regular agitation (150 rpm). Cells were transfected with a plasmid coding for SARS-CoV-2 S RBD using ExpiFectamine 293 transfection reagent, as directed by the manufacturer (Invitrogen) (Beaudoin-Bussieres et al., 2020; Prevost et al., 2020). One week later, cells were pelleted and discarded. Supernatants were filtered using a 0.22 μm filter (Thermo Fisher Scientific). The recombinant RBD proteins were purified by nickel affinity columns, as directed by the manufacturer (Thermo Fisher Scientific). The RBD preparations were dialyzed against phosphate-buffered saline (PBS) and stored in aliquots at −80°C until further use. To assess purity, recombinant proteins were loaded on SDS-PAGE gels and stained with Coomassie Blue.

#### RBD-specific IgG levels and avidity measured by enzyme-Linked Immunosorbent Assay (ELISA)

The SARS-CoV-2 RBD ELISA assay was used to measure the level of RBD-specific IgG, as previously described (Beaudoin-Bussieres et al., 2020; Prevost et al., 2020). Briefly, recombinant SARS-CoV-2 RBD protein was prepared in PBS (2.5 μg/ml) and adsorbed to plates overnight at 4°C. Coated wells were subsequently blocked with blocking buffer then washed. CR3022 monoclonal Ab (50ng/ml) at 1/250, 1/500, 1/1250, 1/2500, 1/5000, 1/10000, 1/20000 dilutions of plasma from SARS-CoV-2-naive or previously infected donors were prepared in a diluted solution of blocking buffer and incubated with the RBD-coated wells. Plates were washed followed by incubation with the respective secondary Abs. Area Under the Cure (AUC) was calculated by using GraphPad. To calculate the RBD-avidity index, we performed a stringent ELISA where the plate was washed with washing buffer supplemented 8M urea. The binding of CR3022 IgG and plasma was quantified with HRP-conjugated antibodies specific for the Fc region of human IgG. HRP enzyme activity was determined after the addition of a 1:1 mix of Western Lightning oxidizing and luminol reagents (Perkin Elmer Life Sciences). Light emission was measured with a LB942 TriStar luminometer (Berthold Technologies).

#### Spike IgG levels measured by cell-based ELISA (CBE)

Detection of the trimeric SARS-CoV-2 S at the surface of HOS cells was performed by a previously described cell-based enzyme-linked immunosorbent assay (ELISA) (Anand et al., 2021). Briefly, parental HOS cells or HOS-Spike cells by Spike specific IgG were seeded in 96-well plates (6×10^4^ cells per well) overnight. Cells were blocked with blocking buffer (10 mg/ml nonfat dry milk, 1.8 mM CaCl_2_, 1 mM MgCl_2_, 25 mM Tris [pH 7.5], and 140 mM NaCl) for 30 min. CR3022 mAb (1 μg/ml) or plasma (at a dilution of 1/250) were prepared in blocking buffer and incubated with the cells for 1h at room temperature. Respective HRP-conjugated anti-human IgG Fc secondary Abs were then incubated with the samples for 45 min at room temperature. For all conditions, cells were washed 6 times with blocking buffer and 6 times with washing buffer (1.8 mM CaCl_2_, 1 mM MgCl_2_, 25 mM Tris [pH 7.5], and 140 mM NaCl). HRP enzyme activity was determined after the addition of a 1:1 mix of Western Lightning oxidizing and luminol reagents (PerkinElmer Life Sciences). Light emission was measured with an LB942 TriStar luminometer (Berthold Technologies). Signal obtained with parental HOS was subtracted for each plasma and was then normalized to the signal obtained with CR3022 mAb present in each plate. The seropositivity threshold was established using the following formula: mean of all SARS-CoV-2 negative plasma + (3 standard deviation of the mean of all SARS-CoV-2 negative plasma).

#### ADCC assay

The SARS-CoV-2 ADCC assay used was previously described (Anand et al., 2021; Beaudoin-Bussieres et al., 2020; Prevost et al., 2020). Briefly, parental CEM.NKr CCR5+ cells were mixed at a 1:1 ratio with CEM.NKr.Spike cells and were stained for viability (Aquavivid: Thermo Fisher Scientific) and a cellular dye (cell proliferation dye eFluor670; Thermo Fisher Scientific) to be used as target cells. Overnight rested PBMCs were stained with another cellular dye (cell proliferation dye eFluor450; Thermo Fisher Scientific), then used as effector cells. Stained target and effector cells were mixed at a ratio of 1:10 in 96-well V-bottom plates. Plasma from SARS-CoV-2 naïve or PI individuals (1/500 dilution) or monoclonal antibody CR3022 (1 μg/mL) were added to the appropriate wells. The plates were subsequently centrifuged and incubated at 37°C, 5% CO_2_ for 5 hours before being fixed in a 2% PBS-formaldehyde solution. All samples were acquired on an LSRII cytometer (BD Biosciences) and data analysis was performed using FlowJo v10.7.1 (Tree Star).

#### Virus neutralization assay

The SARS-CoV-2 virus neutralization assay used was previously (Prevost et al., 2020). Briefly, 293T cells were transfected with the lentiviral vector pNL4.3 R-E-Luc plasmid (NIH AIDS Reagent Program) and a plasmid encoding for the full-length SARS-CoV-2 Spike D614G glycoprotein (Beaudoin-Bussieres et al., 2020; Prevost et al., 2020) at a ratio of 10:1. Two days post-transfection, cell supernatants were harvested and stored at −80°C until use. Pseudoviral particles were incubated with the indicated plasma dilutions (1/50; 1/250; 1/1250; 1/6250; 1/31250) for 1h at 37°C and were then added to the 293T-ACE2 target cells followed by incubation for 48h at 37°C. Then, cells were lysed and followed by one freeze-thaw cycle. An LB942 TriStar luminometer (Berthold Technologies) was used to measure the luciferase activity. The neutralization half-maximal inhibitory dilution (ID50) represents the plasma dilution to inhibit 50% of the infection of 293T-ACE2 cells by SARS-CoV-2 pseudoviruses.

#### SARS-CoV-2-specific B cells characterization

To detect SARS-CoV-2-specific B cells, we conjugated recombinant RBD proteins with Alexa Fluor 488 or Alexa Fluor 594 (Thermo Fisher Scientific) according to the manufacturer’s protocol. 2 × 10^6^ frozen PBMC from SARS-CoV-2 naïve and previously-infected donors were prepared in Falcon^®^ 5ml-round bottom polystyrene tubes at a final concentration of 4 × 10^6^ cells/mL in RPMI 1640 medium (GIBCO) supplemented with 10% of fetal bovine serum (Seradigm), Penicillin-Streptomycin (GIBCO) and HEPES (GIBCO). After a rest of 2h at 37°C and 5% CO_2_, cells were stained using Aquavivid viability marker (GIBCO) in DPBS (GIBCO) at 4°C for 20min. The detection of SARS-CoV-2-antigen specific B cells was done by adding the RBD probes to the antibody cocktail listed in Table S1. Staining was performed at 4°C for 30min and cells were fixed using 2% paraformaldehyde at 4°C for 15min. Stained PBMC samples were acquired on Symphony cytometer (BD Biosciences) and analyzed using FlowJo v10.8.0 software.

#### Activation-induced marker (AIM) assay

The AIM assay was adapted for SARS-CoV-2 specific CD4 and CD8 T cells was as previously described (Tauzin et al., 2021b). PBMCs were thawed and rested for 3h in 96-well flat-bottom plates in RPMI 1640 supplemented with HEPES, penicillin and streptomycin and 10% FBS. 1.7×10^6^ PBMCs were stimulated with a S glycoprotein peptide pool (0.5 μg/ml per peptide, corresponding to the pool of 315 overlapping peptides (15-mers) spanning the complete amino acid sequence of the Spike glycoprotein (JPT) for 15h at 37 °C and 5% CO_2_. CXCR3, CCR6, CXCR6 and CXCR5 antibodies were added in culture 15 min before stimulation. A DMSO-treated condition served as a negative control and *Staphylococcus enterotoxin B* SEB-treated condition (0.5 μg/ml) as positive control. Cells were stained for viability dye for 20 min at 4 °C then surface markers (30 min, 4 °C). Abs used are listed in the Table S2. Cells were fixed using 2% paraformaldehyde for 15min at 4°C before acquisition on Symphony cytometer (BD Biosciences). Analyses were performed using FlowJo v10.8.0 software.

#### Intracellular Cytokine Staining (ICS)

The ICS assay adapted to study SARS-CoV-2-specific T cells was previously described (Tauzin et al., 2021b). PBMCs were thawed and rested for 2 h in RPMI 1640 medium supplemented with 10% FBS, Penicillin-Streptomycin (Thermo Fisher scientific, Waltham, MA) and HEPES (Thermo Fisher scientific, Waltham, MA). 1.7×10^6^ PBMCs were stimulated with a S glycoprotein peptide pool (0.5 μg/mL per peptide from JPT, Berlin, Germany) corresponding to the pool of 315 overlapping peptides (15-mers) spanning the complete amino acid sequence of the S glycoprotein.

Cell stimulation was carried out for 6h in the presence of mouse anti-human CD107a, Brefeldin A and monensin (BD Biosciences, San Jose, CA) at 37 °C and 5% CO_2_. DMSO-treated cells served as a negative control, and SEB as positive control. Cells were stained for Aquavivid viability marker (Thermo Fisher scientific, Waltham, MA) for 20 min at 4 °C and surface markers (30 min, 4 °C), followed by intracellular detection of cytokines using the IC Fixation/Permeabilization kit (Thermo Fisher scientific, Waltham, MA) according to the manufacturer’s protocol before acquisition on a Symphony flow cytometer (BD Biosciences) and analysis using FlowJo v10.8.0 software. Abs used are listed in the Table S3.

### QUANTIFICATION AND STATISTICAL ANALYSIS

#### Statistical analysis

Symbols represent biologically independent samples from SARS-CoV-2 naïve individuals and SARS-CoV-2 PI individuals. Lines connect data from the same donor. Statistics were analyzed using GraphPad Prism version 8.4.3 (GraphPad, San Diego, CA). Every dataset was tested for statistical normality and this information was used to apply the appropriate (parametric or nonparametric) statistical test. Differences in responses for the same patient before and after vaccination were performed using Wilcoxon matched pair tests. Differences in responses between naïve and PI individuals were measured by Mann-Whitney tests. P values <0.05 were considered significant. P values are indicated for each comparison assessed. For correlations, Spearman’s R correlation coefficient was applied. For graphical representation on a log scale (but not for statistical tests), null values were arbitrarily set at the minimal values for each assay.

#### Software scripts and visualization

Graphics and pie charts were generated using GraphPad PRISM version 8.4.1 and ggplot2 (v3.3.3) in R (v4.1.0). Heat maps were generated in R (v4.1.0) using the *pheatmap* package (v1.0.12). Principal component analyses were performed with the *prcomp* function (R). Uniform manifold approximation and projection (UMAP) was performed using package M3C (v1.14.0) on gated FCS files loaded through the flowCore package (v2.4.0). Samples were downsampled to a comparable number of events (300 cells for AIM, 100 cells for ICS). Scaling and logicle transformation of the flow cytometry data was applied using the FlowSOM (Quintelier et al., 2021) R package (v2.0.0). All samples from naïve and PI at all time points were loaded. Clustering was achieved using Phenograph (v0.99.1) with the hyperparameter k (number of nearest-neighbors) set to 150). R codes scripted for this paper is provided as supplementary material. We obtained an initial 15 AIM^+^ and 11 cyto^+^ clusters. After careful examination, five low-abundance AIM^+^ clusters were merged based on proximity on the UMAP, phenotypic similarities and concomitant longitudinal trajectories. This resulted in a final 10 AIM+ clusters. None of the 11 cyto+ clusters were merged. For B and CD4^+^ T cell phenotyping, only participants with >5 RBD+ B events across all depicted time points were analyzed.

## KEY RESOURCES TABLE

| REAGENT or RESOURCE | SOURCE | IDENTIFIER |
| --- | --- | --- |
| **Antibodies** |  |  |
| UCHT1 (BUV395) [Human anti-CD3] | BD Biosciences | Cat#563546 ;<br>Lot:9058566 ;<br>RRID:AB_2744387 |
| UCHT1 (BUV496) [Human anti-CD3] | BD Biosciences | Cat#612941 ;<br>Lot:1022424 ;<br>RRID:AB_2870222 |
| L200 (BV711) [Human anti-CD4] | BD Biosciences | Cat#563913 ;<br>Lot:03000025;<br>RRID:AB_2738484 |
| SK3 (BB630) [Human anti-CD4] | BD Biosciences | Cat#624294<br>CUSTOM ;<br>Lot:0289566 |
| RPA-T8 (BV570) [Human anti-CD8] | Biolegend | Cat#301037 ;<br>Lot:B281322 ;<br>RRID:AB_10933259 |
| M5E2 (BUV805) [Human anti-CD14] | BD Biosciences | Cat#612902 ;<br>Lot:0262150 ;<br>RRID:AB_2870189 |
| M5E2 (BV480) [Human anti-CD14] | BD Biosciences | Cat#746304; Lot :<br>9133961 ;<br>RRID:AB_2743629 |
| 3G8 (BV650) [Human anti-CD16] | Biolegend | Cat#302042 ;<br>Lot:B323847 ;<br>RRID:AB_2563801 |
| HIB19 (APC-eFluor780) [Human anti-CD19] | Thermo Fisher Scientific | Cat#47-0199 ;<br>Lot:2145095 ;<br>RRID:AB_1582231 |
| HIB19 (BV480) [Human anti-CD19] | BD Biosciences | Cat#746457 ;<br>Lot:1021649 ;<br>RRID:AB_2743759 |
| HI100 (PerCP Cy5.5) [Human anti-CD45RA] | BD Biosciences | Cat#563429 ;<br>Lot:8332746 ;<br>RRID:AB_2738199 |
| NCAM16.2 (BUV737) [Human anti-CD56] | BD Biosciences | Cat#564448 ;<br>Lot:8288818 ;<br>RRID:AB_2744432 |
| FN50 (PerCP-eFluor710) [Human anti-CD69] | Thermo Fisher Scientific | Cat#46-0699-42 ;<br>Lot:1920361 ;<br>RRID:AB_2573694 |
| FN50 (BV650) [Human anti-CD69] | Biolegend | Cat# 310934 ;<br>Lot:B303462 ;<br>RRID:AB_2563158 |
| H4A3 (BV786) [Human anti-CD107A] | BD Biosciences | Cat#563869 ;<br>Lot:8144866 ;<br>RRID:AB_2738458 |
| ACT35 (APC) [Human anti-CD134 (OX40)] | BD Biosciences | Cat#563473 ;<br>Lot:1015537 ;<br>RRID:AB_2738230 |
| 4B4-1 (PE-Dazzle 594) [Human anti-CD137 (4-1BB)] | Biolegend | Cat# 309826 ;<br>Lot:B253152 ;<br>RRID:AB_2566260 |
| TRAP1 (BV421) [Human anti-CD154 (CD40L)] | BD Biosciences | Cat#563886 ;<br>Lot:9037850 ;<br>RRID:AB_2738466 |
| TRAP1 (PE) [Human anti-CD154 (CD40L)] | BD Biosciences | Cat#555700 ;<br>Lot:7086896 ;<br>RRID:AB_396050 |
| J25D4 (BV421) [Human anti-CD185 (CXCR5)] | Biolegend | Cat# 356920 ;<br>Lot:B325837 ;<br>RRID:AB_2562303 |
| B27 (PECy7) [Human anti-IFN-γ] | BD Biosciences | Cat#557643 ;<br>Lot:8256597 ;<br>RRID:AB_396760 |
| MQ1-17H12 (PE-Dazzle594) [Human anti-IL-2] | Biolegend | Cat#500344 ;<br>Lot:B2261476 ;<br>RRID:AB_2564091 |
| JES3-9D7 (PE) [Human anti-IL-10] | BD Biosciences | Cat#554498 ;<br>Lot:8198773 ;<br>RRID:AB_395434 |
| eBio64CAP17 (eFluor660) [Human anti-IL-17A] | Thermo Fisher Scientific | Cat#50-7179-42 ;<br>Lot:2151998 ;<br>RRID:AB_11149126 |
| Mab11 (Alexa Fluor 488) [Human anti-TNF-α] | Biolegend | Cat#502915 ;<br>Lot:B285221 ;<br>RRID:AB_493121 |
| LIVE/DEAD Fixable dead cell | Thermo Fisher Scientific | L34960 |
| Biological samples |  |  |
| SARS-CoV-2 naïve donor blood samples |  |  |
| SARS-CoV-2 prior infection donor blood samples |  |  |
| Chemicals, peptides, and recombinant proteins |  |  |
| PepMix™ SARS-CoV-2 (Spike Glycoprotein) | JPT | Cat#PM-WCPV-S-1 |
| Staphylococcal Enterotoxin B (SEB) | Toxin technology | Cat#BT202 |
| Experimental models: cell lines |  |  |
| Software and algorithms |  |  |
| Flow Jo v10.8.0 | Flow Jo | https://www.flowjo.com |
| GraphPad Prism v8.4.1 | GraphPad | https://www.graphpad.com |
| R studio v4.1.0 | R studio | https://rstudio.com |

