## Supplemental Text and Figures for "Temporal associations of B and T cell immunity with robust vaccine responsiveness in a 16-week interval BNT162b2 regimen"

Supplemental Text and Figures Click here to access/download;Supplemental Text and Figures;Nayrac et al Supplementary text and figures.pdf



#### SUPPLEMENTARY FIGURE LEGENDS

**Figure S1. Related to figure 1. (A)** Gating strategy to identify RBD-specific B cell responses. **(B)** Comparisons between RBD-B cell responses at the V1 timepoint for PI and at the V3 timepoint for naïve participants. A Mann-Whitney test is indicated above the graph. Data represented naïve (blue; n=22) and SARS-CoV-2 pre-infected (PI) individuals (orange, n=21). **(C)** Longitudinal RBD-specific B cell responses in pre-infected (PI) participants that did not receive a second dose after V2 (n=12). Lines connect data from the same donor. The bold line represents the median value of each cohort. Wilcoxon tests for each pairwise comparison are shown underneath. **(D)** Examples of gatings for IgD, IgM, IgA and IgG expression on total CD19$^{+}$CD20$^{+}$ B cells or RBD-specific B cells. **(E)** Histograms quantifying the frequency of IgD$^{+}$, IgM$^{+}$, IgA$^{+}$ and IgG$^{+}$ RBD-specific B cells at different timepoints, comparing naïve (n=22) vs PI participants (n=11). Mann-Whitney test are shown. **(F)** Longitudinal trajectories of isotype expression frequencies in naïve (n=22) and PI (n=11) participants. Lines connect data points for individual participants. Wilcoxon tests are shown above each panel. **(G)** Example of the gating strategy of IgD and CD27 co-expression on RBD-specific B cells. In support to the pie charts displayed in Figure 1G. **(H)** Histograms showing the proportion of IgG+ in IgD-CD27- RBD+ B cells. Mann-Whitney test are shown above. **(I)** Histograms reporting the longitudinal frequency of each IgD and CD27 RBD-B phenotypes in CD19+CD20+ B cells for naïve (n=22) and PI (n=11) participants. In support to the pie charts displayed in Figure 1F. Wilcoxon tests are shown above.

**Figure S2: Related to figure 2. (A)** Representative upstream generic gating and **(B)** ORgate strategy to identify SARS-CoV-2-specific AIM$^{+}$ T cells. For simplicity, the example focuses on CD4$^{+}$ T cells. **(CD)** Raw frequencies of **(C)** AIM$^{+}$ CD4+ and **(D)** CD8$^{+}$ T cells following *ex vivo* stimulation of PBMCs with a pool of SARS-CoV-2 Spike peptides. As a control, PBMCs cells were left unstimulated (grey bars). The data for naïve (blue, n=26) and pre-infected (PI; orange, n=27) individuals are displayed. The bars represent median values. Mann-Whitney tests are shown. The number of conditions reaching >2x no Ag are shown below each timepoint. **(E)** Representative ORgate strategy to identify SARS-CoV-2-specific cytokine-expressing T cells. For simplicity, the example focuses on CD4 T cells. **(F)** Raw frequencies of cytokine-expressing $CD4^+$ T cells following *ex vivo* stimulation of PBMCs with a pool of SARS-CoV-2 Spike peptides. As a control, PBMCs cells were left unstimulated (grey bars). The data for naïve (blue, n=26) and pre-infected (PI; orange, n=27) individuals are displayed. The bars represent median values. Mann-Whitney tests are shown. The number of conditions reaching >2x no Ag are shown below each time point. **(G)** Comparisons between $AIM^+CD4^+$ (red) and $AIM^+CD8^+$ (purple) T cell responses. Median and interquartile range are shown, with Mann-Whitney tests. **(H)** Raw frequencies of cytokine-expressing $CD8^+$ T cells following *ex vivo* stimulation of PBMCs with a pool of SARS-CoV-2 Spike peptides. As a control, PBMCs cells were left unstimulated (grey bars). The data for naïve (blue, n=26) and pre-infected (PI; orange, n=27) individuals are displayed. The bars represent median values. Mann-Whitney tests are shown. The number of conditions reaching >2x no Ag are shown below each timepoint. **(IJ)** Longitudinal $AIM^+CD4^+$ (**I**), $CD8^+$ (**J**) and $cytokine^+$ $CD4^+$ T cell responses in pre-infected (PI) participants that did not receive a second dose after V2 (n=12). Lines connect data from the same donor. The bold line represents the median value of each cohort. Wilcoxon test for each pairwise comparison is shown underneath.

**Figure S3: Related to figure 3. (A-C)** Multivariate analysis. **(A)** Heat map overlaid on the $AIM^+$ UMAP showing the gradient of expression for each marker. **(B)** The longitudinal net frequency of $AIM^+$ in $CD4^+$ T cells for clusters 4, 7, 8 and 9 for naïve (blue, n=22) and PI (orange; n=11) participants. Wilcoxon tests are shown beside for each pairwise comparison. Complement Figure 3D. **(C)** Cohort comparisons, with Mann-Whitney tests. **(C-F)** Univariate analyses. Example of (**C**) $CXCR5^+$, (**D**) $CXCR3^+$, (**E**) $CXCR6^+$ and (**F**) $CCR6^+$ gating on total and $AIM^+$ populations for univariate analyses. (**HI**) Net frequencies of AIM$^+$CCR6$^+$CD4$^+$ T cells. (**H**) Longitudinal analysis presenting both naïve and PI are overlaid. Wilcoxon tests are shown besides each panel. Lines connect data from the same donor. Bold lines represent median values. **(I)** Histogram comparing naïve and PI participants. Mann-Whitney tests are shown.

**Figure S4: Related to Figure 4. (A-C)** Multivariate analysis. **(A)** Heat map overlaid on the cytokine$^+$ UMAP showing the gradient of expression for each marker. **(B)** Longitudinal net frequencies of cytokine$^+$CD4$^+$ T cells for clusters 7, 8, 9, 10 and 11 for naïve (blue, n=22) and PI (orange; n=11) participants. Wilcoxon tests are shown for each pairwise comparison. Support figure 4D. **(C)** Cohort comparison with Mann-Whitney tests.

**Figure S5: Related to Figure 5.** Univariate correlations for validation. (**A**) Correlation between total AIM$^+$CD4$^+$ T cell frequencies at V0-V3 and RBD-specific B cell frequencies at V3 (n=21). (**B**) Correlation between total cytokine$^+$CD4$^+$ T cell frequencies at V0-V3 and AIM$^+$CD8$^+$ T cell frequencies at V3 (n = 19). (**C**) The r and p values from a Spearman test are indicated in each graph.

**Figure S6: Related to Figure 6.** AIM$^+$ (A) and cytokine$^+$ (B) CD4$^+$ T cell sub-PCA analyses. The PC coordinates were set based on the primary PCA combining all timepoints. PC coordinates were then plotted by timepoints for clarity. **(A)** The top panels present the PCA plots. The proportion of the variance attributed to PC1 and PC2 are indicated on the axes. The numbers of participants analyzed in each PCA plot are indicated in each plot. Box and whisker plots of the PC1 and PC2 between group are presented below, with Mann-Whitney tests. In **BC**, blue is representing naïve participants and orange, PI.

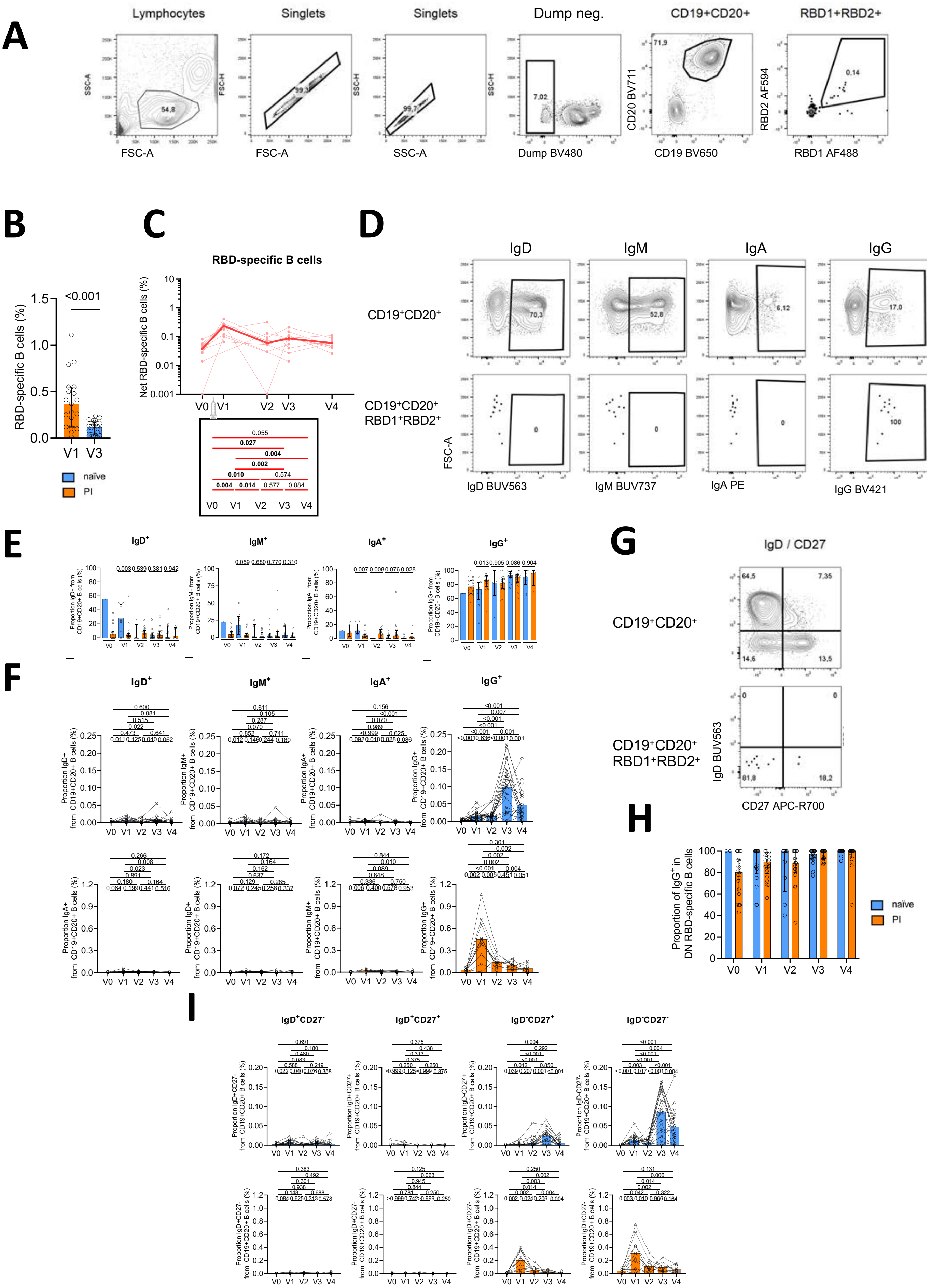


### Supplemental Figure 1

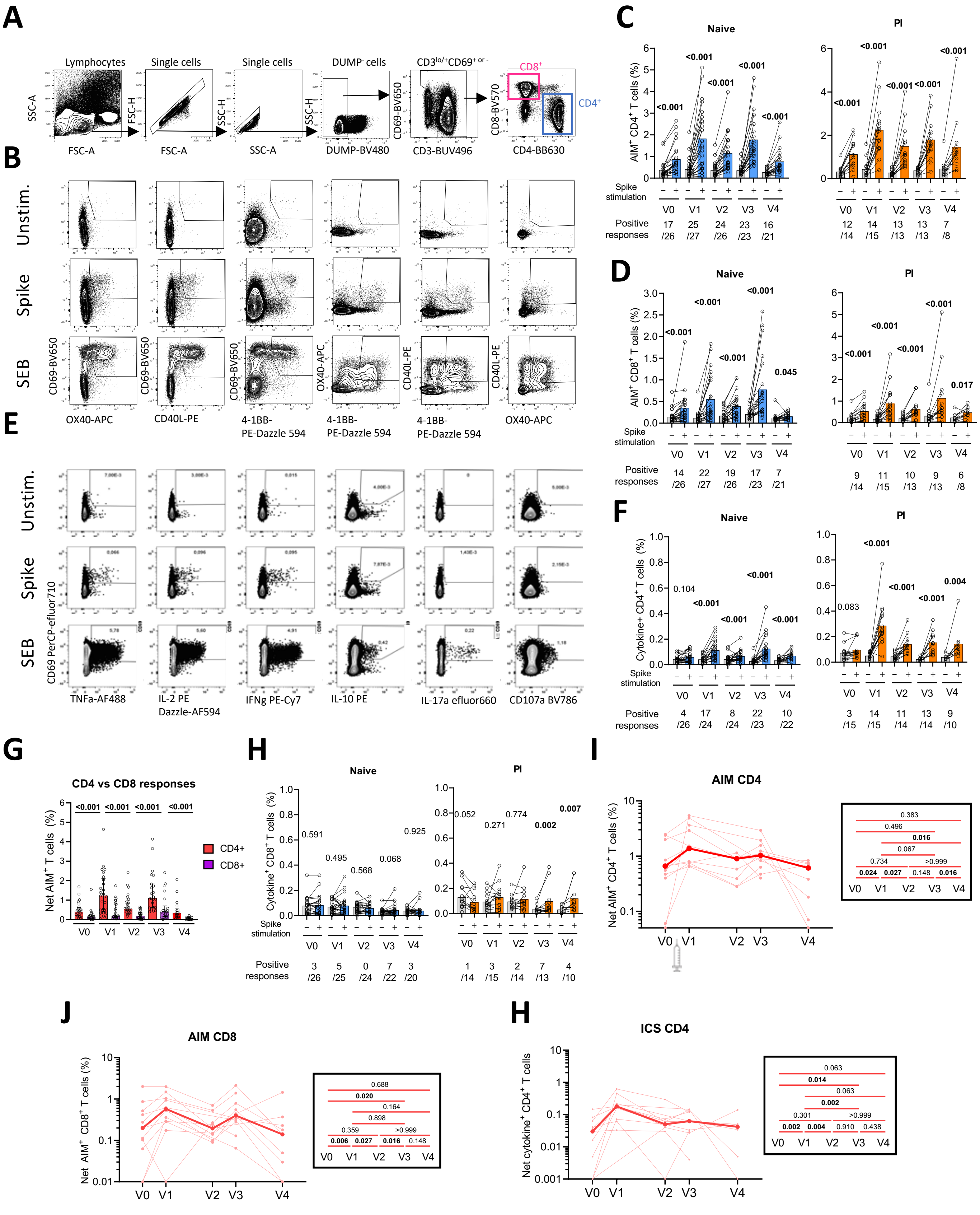


### Supplemental Figure 2

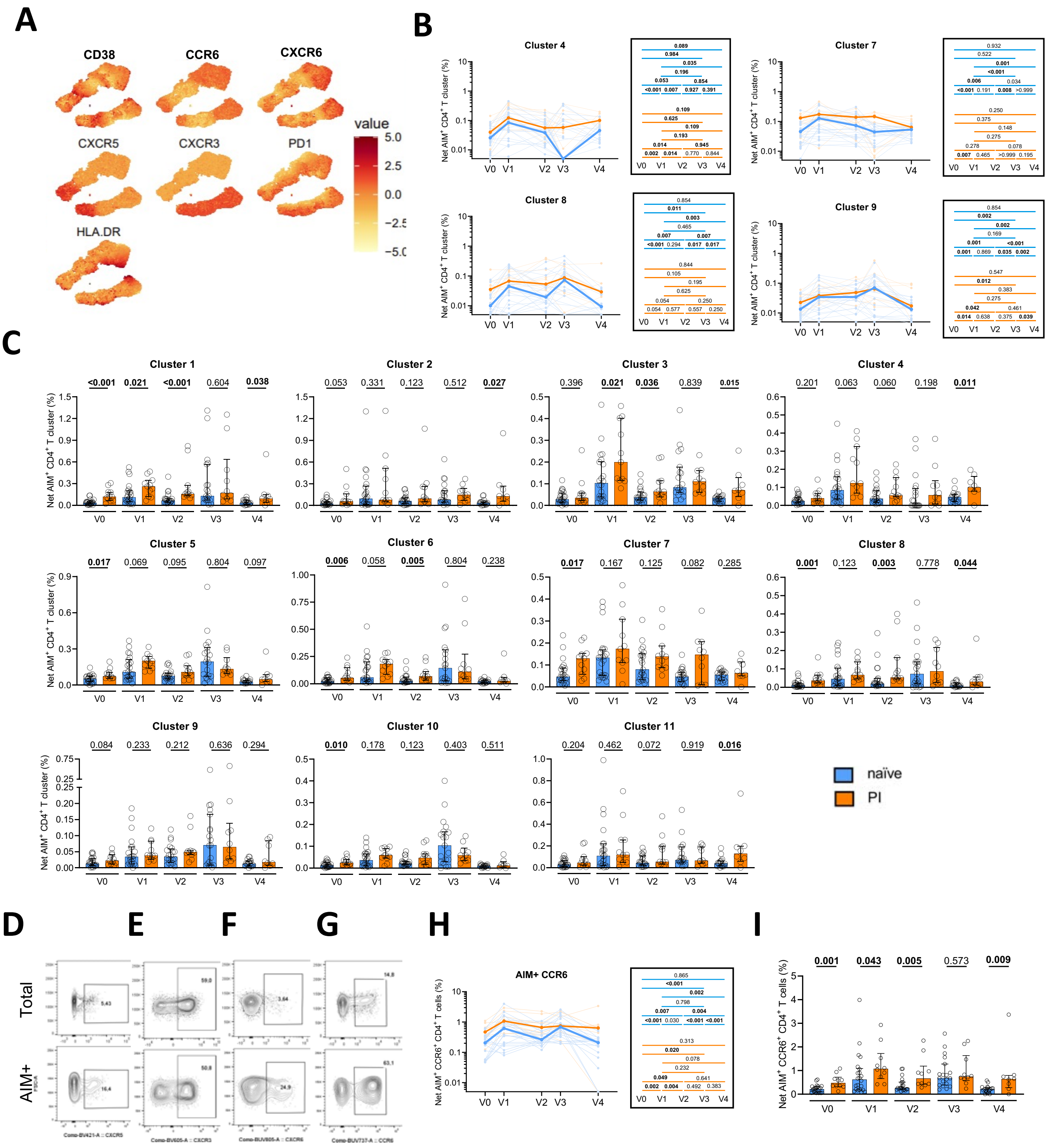


### Supplemental Figure 3

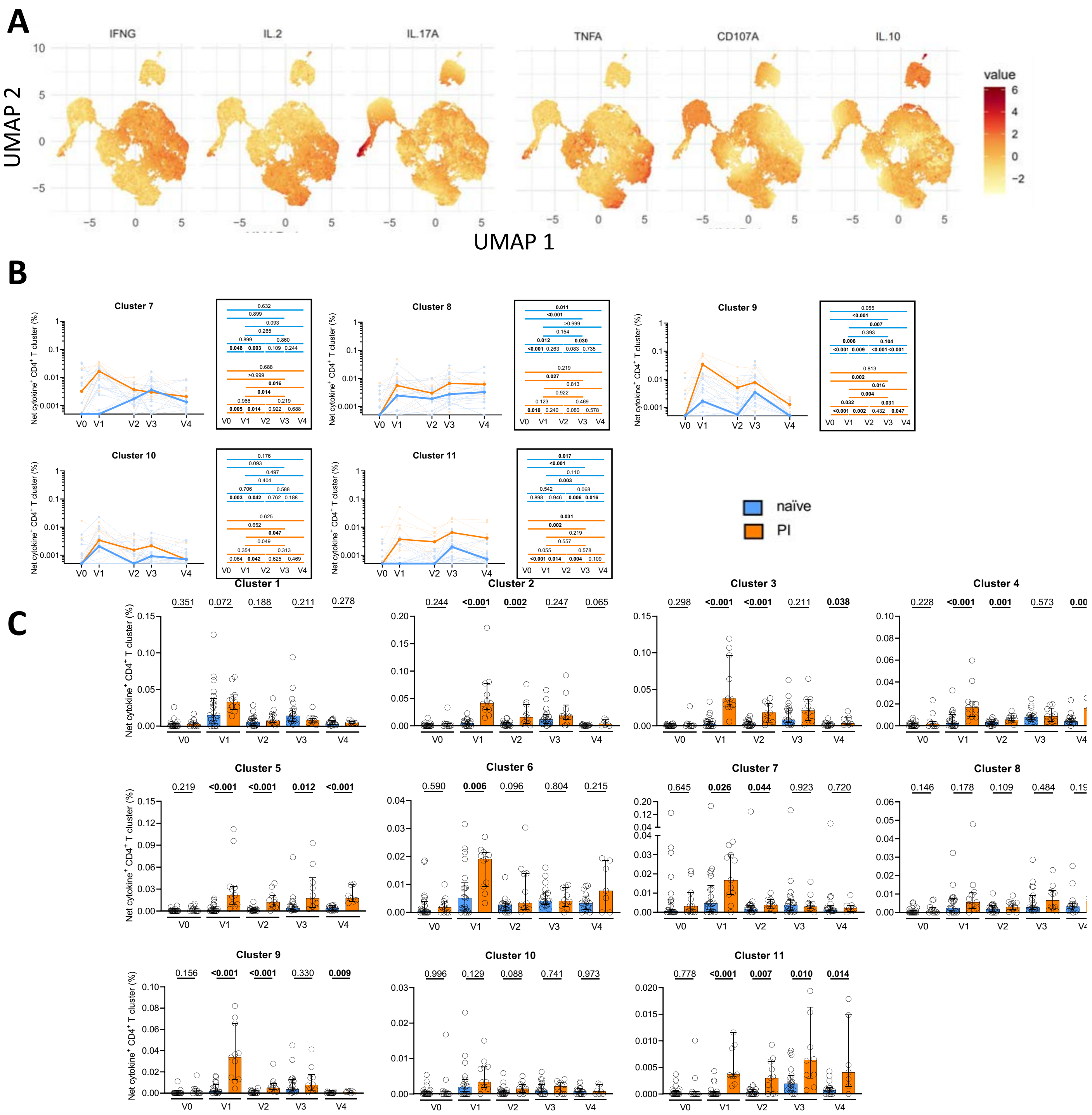


**Supplemental Figure 4**

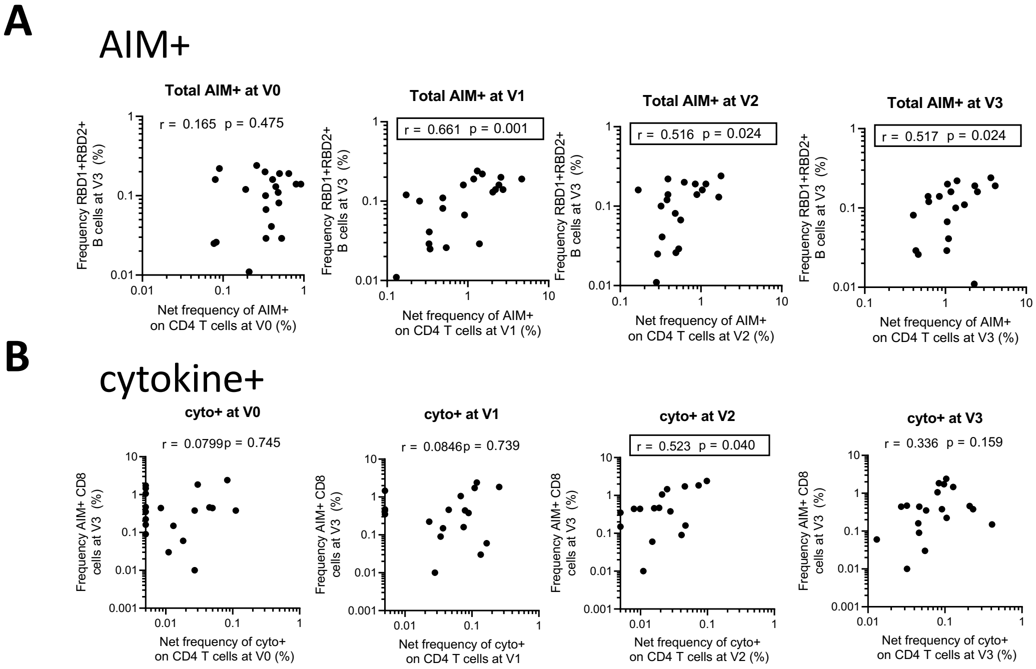


### Supplementary Figure 5

**A** AIM$^+$

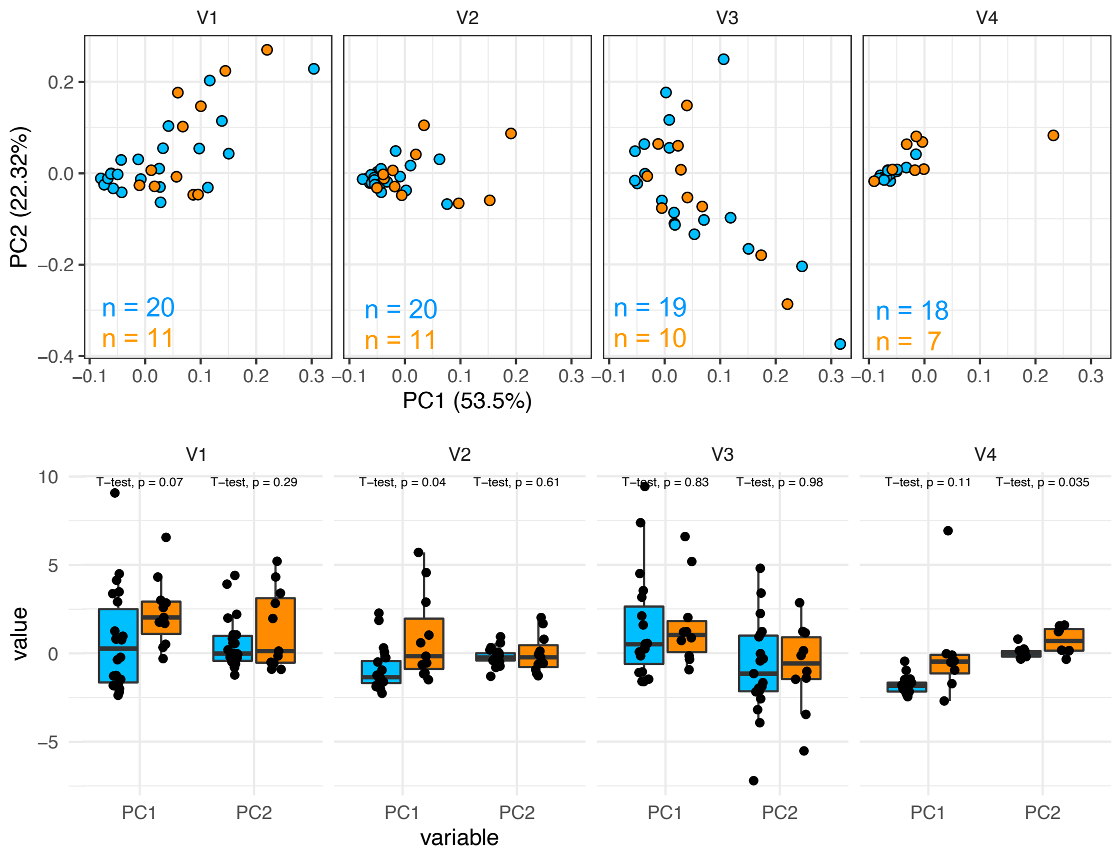


**B** Cytokine$^+$

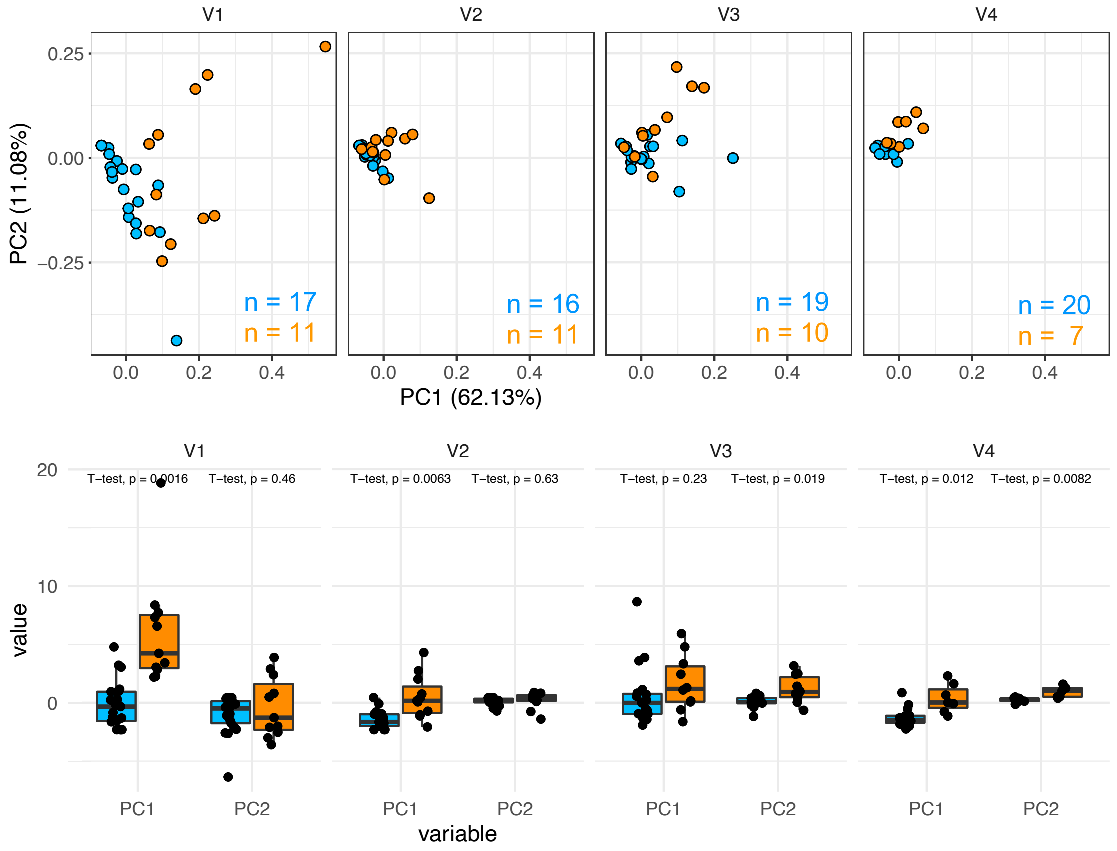


### Supplementary figure 6
